# Frustration Quenching and Network Topology of the Energy Landscape as Primary Determinants of Protein–Ligand Binding Pose Prediction by Deep Learning Models

**DOI:** 10.64898/2026.10.05.756861

**Authors:** Lucas Turano, Brandon Foley, Ryan Dennis, Will Gatlin, Gennady M. Verkhivker

**Affiliations:** Keck Center for Science and Engineering, Department of Biological Sciences, Schmid College of Science and Technology, Chapman University, Orange, CA 92866, United States of America; Department of Biomedical and Pharmaceutical Sciences, Chapman University School of Pharmacy, Irvine, CA 92618, United States of America

## Abstract

Deep learning co-folding and docking models accurately predict ligand binding poses at orthosteric sites yet systematically underperform at allosteric pockets. Here, we demonstrate that this accuracy gap reflects a fundamental biophysical property of local energy landscape topology—specifically, the magnitude of frustration quenching upon ligand binding—rather than an intrinsic algorithmic defect. By evaluating leading co-folding and docking paradigms across curated benchmarks and large-scale uncurated datasets, we show that AI prediction success strongly tracks frustration quenching—the degree to which ligand binding relieves local energetic strain to form a steep energy funnel. Topologically, frustration-weighted residue interaction networks reveal that orthosteric pockets act as high-betweenness communication hubs embedded within contiguous, minimally frustrated pathways. In contrast, allosteric sites reside in low-betweenness peripheral regions with fragmented frustration profiles, presenting flatter, entropy-dominated landscapes that challenge structure prediction algorithms. Comparing generative diffusion with single-shot regression demonstrates that multi-pose sampling capacity is required to cross energy barriers and discover the frustration funnel, whereas joint receptor– ligand coordinate co-adaptation governs local optimization within the funnel. Collectively, this study reframes the allosteric “blind spot” as a quantitative property of energy landscape architecture rather than a defect of any specific predictive framework. We demonstrate that frustration quenching magnitude and structural network centrality serve as physical descriptors that define the intrinsic limits of AI performance, reframing AI accuracy from a benchmark metric into a physics-grounded quantitative reporter of macromolecular binding landscape architecture.

## Introduction

Artificial intelligence (AI) has fundamentally reshaped structural biology. AlphaFold2 near-experimental protein structure prediction^1,2^ marked a watershed moment, followed by extensions to multi-chain assembly^3^, sequence-based protein language models (PLMs) such as ESMFold^4^ and RoseTTAFold^5,6^ and programmable generative frameworks for de novo design.^7–10^ These advances have dramatically accelerated structure-based drug discovery and opened new routes to engineering proteins with tailored binding properties. Despite this progress, allosteric regulation remains a fundamental blind spot for current AI models.^11–14^ Allostery is intrinsically an ensemble phenomenon requiring proteins to interconvert between conformational substates whose relative populations are modulated by ligand binding.^11–13^ Contemporary structure predictors, by contrast, have been optimized to recover single thermodynamically dominant ground states and consequently struggle to sample the conformational heterogeneity on which allosteric mechanisms depend.^15–20^ Post hoc remedies such as MSA subsampling, in silico mutagenesis, and clustering-based sampling have yielded only incremental improvements, limited by the evolutionary priors that govern training and by the sparsity of allosteric examples in structural databases.^16–20^

Concurrently, biomolecular binding prediction has undergone a paradigm shift from classical physics-based docking toward end-to-end AI co-folding. Diffusion-based models now jointly generate protein conformations and ligand poses. DiffDock reframed docking as generative sampling over the non-Euclidean manifold of ligand poses^21^ with a refined successor improving generalization.^22^ AlphaFold3 (AF3) unified folding and docking by diffusing an atomic “cloud” of protein and ligand coordinates into a synchronized holo state,^23^ outperforming classical tools such as AutoDock Vina^24^ by a wide margin on the PoseBusters benchmark.^25^ Open-source reproductions such as Protenix^26^ have achieved comparable accuracy, while Boltz-2 augments co-folding with affinity and binder-classification heads trained on experimental and MD-derived data.^27^ Chai-1 fuses PLM embeddings with paired attention for high-fidelity predictions from sparse evolutionary signal^28^ and DynamicBind explicitly models ligand-induced receptor rearrangements via an SE(3)-equivariant diffusion framework, enabling cryptic pocket formation and large-scale conformational morphing.^29^ Complementary architectures operate without co-folding: EquiBind predicts binding poses in a single shot from apo structures,^30^ DiffSBDD generates structure-based ligands via equivariant diffusion^31^ and TANKBind segments proteins into functional blocks for trigonometry-aware prediction.^32^ Together, these models constitute a rich and diverse methodological landscape, yet all share a common performance profile when confronted with allosteric sites.

In parallel, the prediction of druggable binding sites has expanded into a broad ecosystem spanning geometric cavity detectors (Fpocket^33^, SiteHound^34^ and MetaPocket^35^), machine learning tools (P2Rank^36^), deep learning architectures (DeepSite^37^), PLM-based annotators^38,39^ and physics-informed methods AlloPred,^40^ Allosite,^41^ AllositePro,^42^ and DeepAllo^43^. Cryptic pocket predictors such as PocketMiner^44^ have extended this landscape to transient sites. Across this ecosystem, however, a consistent pattern emerges where methods perform well on orthosteric sites but exhibit a pronounced and systematic drop in accuracy at allosteric and cryptic sites.^45,46^ This limitation suggests that the difficulty of allosteric prediction may not be solely algorithmic but may instead reflect a fundamental property of the underlying binding landscapes.

We have initially proposed that this pervasive underperformance reflects a fundamental biophysical mismatch between the energy landscapes of orthosteric and allosteric binding sites.^47^ Orthosteric pockets are typically defined by strong evolutionary conservation, geometric recurrence, and pronounced energetic stabilization—features that create steep, minimally frustrated funnels that tightly align with the inductive biases of contemporary AI architectures.^48–51^ Allosteric sites, by contrast, frequently reside in evolutionarily permissive and conformationally adaptive regions.^52–54^ Because their functional versatility relies on neutral frustration and structural plasticity rather than dominant, recurrent motifs, these sites lack the explicit geometric and energetic signatures that pattern-recognition models depend on for pose convergence.^47,55–57^ Local frustration analysis offers a particularly incisive lens for interrogating this hypothesis.^48–54^ By comparing the energetics of native residue–residue contacts against decoy ensembles, frustration analysis partitions proteins into minimally frustrated regions optimized for stability, highly frustrated regions associated with strain or functional strain, and neutrally frustrated regions compatible with conformational plasticity. While the FrustratometeR package^58^ enabled structured calculations across point mutants and molecular dynamics trajectories, traditional physics-based frustration calculations scale quadratically with protein length and require evaluating thousands of mutated structural variants per residue—creating a severe computational bottleneck that renders proteome-scale studies impractical. To overcome this limitation, scalable deep learning tools have recently emerged. FrustraMPNN leverages transfer learning from ProteinMPNN with a message-passing graph architecture and light attention to predict single-residue local frustration indices directly from 3D backbone coordinates at ultra-fast speeds.^59^ Bypassing structural input requirements entirely, FrustrAI-Seq fine-tunes ProtT5 protein language model embeddings via low-rank adaptation (LoRA) to infer residue-level frustration directly from amino acid sequence alone—delivering a three-order-of-magnitude speedup capable of profiling the human proteome in minutes.^60^ In the context of AI binding prediction, frustration provides an interpretable biophysical readout: orthosteric ligand binding quenches local frustration to enforce tight geometric convergence, whereas allosteric sites retain neutral frustration that offers no directional signal to guide AI models.^47,51,55–57^

In our previous study, we developed a dual explainable AI framework that combined rigorous benchmarking of five co-folding models on stratified orthosteric and allosteric datasets with physics-based frustration analysis.^47^ That work established a universal allosteric blind spot across AF3, Protenix, Boltz-2, Chai-1, and DynamicBind and demonstrated that the dichotomy in prediction performance is not algorithmic in origin but is encoded in the energetic grammar of the underlying binding landscapes. Orthosteric sites undergo ligand-induced frustration quenching, transitioning from strained apo states to minimally frustrated holo states and creating steep funnels that enforce convergence. Allosteric sites remain neutrally frustrated throughout, permitting multiple geometrically distinct solutions with comparable interaction topologies. That study further revealed a systematic decoupling between contact topology recovery (QS-score remains moderately high) and geometric precision (ligand RMSD doubles), indicating that models recover the vocabulary of allosteric interactions but fail to resolve their syntax.^47^

However, several critical questions remained unresolved. First, it was unclear whether prediction accuracy consistently tracks frustration quenching magnitude across the full spectrum of binding site architectures. Second, because prior evaluations focused exclusively on co-folding models that jointly predict protein and ligand coordinates, whether this frustration–predictability relationship is paradigm-independent—extending to classical docking models that treat the protein as a rigid or semi-rigid receptor—remained untested.

To address these gaps, we expand the analysis in four directions. First, we incorporate the MISATO dataset^61^ as a general-purpose benchmark. MISATO is derived from PDBbind without curation for binding site type, providing an unbiased sampling of protein–ligand interfaces across diverse protein families, ligand chemotypes, and binding site architectures. Because it contains a mixture of orthosteric-like, intermediate, and allosteric-like complexes, MISATO enables direct testing of whether the prediction accuracy tracks frustration quenching magnitude across a broad spectrum of binding sites. Second, we add two docking models that occupy opposite extremes of the docking paradigm: DiffDock^21^ a generative diffusion model that samples multiple poses and ranks them by confidence, and EquiBind^30^ a rigid regression model that predicts a single pose in one step without refinement. Comparing these architectures allows us to probe whether sampling capacity determines access to the frustration funnel. We expand our evaluation metrics to include LDDT-PLI, which assesses the preservation of local atomic distances and side-chain interactions within the binding pocket. This metric bridges global topology (QS-score) and global geometry (RMSD), testing whether topology–geometry decoupling extends to local pocket interactions. We perform comprehensive frustration and Protein Structure Network (PSN) analyses across all datasets to determine whether topology–frustration coupling represents a universal property of protein–ligand recognition.

Collectively, this study reframes the allosteric “blind spot” as a quantitative property of energy landscape architecture rather than a defect of any specific predictive framework. By extending our evaluation to an uncurated benchmark and two paradigmatically opposed docking models, we demonstrate that frustration quenching magnitude serves as a model-independent predictor of prediction success. Furthermore, we show that this same energetic variable organizes both the spatial distribution of frustration within the protein and the communication networks that connect the binding site to the rest of the fold—findings with direct implications for benchmarking, model design, and the identification of tractable allosteric sites for AI-guided drug discovery.

## Materials and Methods

### Datasets

This study was designed to isolate the physical determinants of AI co-folding performance through controlled comparison of orthosteric and allosteric binding across rigorously curated structural datasets, complemented by a broad, general-purpose benchmark. Five state-of-the-art co-folding architectures (AF3^23^, Protenix,^26^ Boltz-2,^27^ Chai-1^28^ and DynamicBind^29^) were evaluated on stratified orthosteric and allosteric datasets using metrics that distinguish geometric accuracy from contact topology recovery. Two additional docking methods (DiffDock^21^ and EquiBind^30^) were evaluated on the MISATO dataset^61^ to enable direct comparison between co-folding and docking paradigms. Local frustration analysis^48–54^ was used to link prediction outcomes to the underlying energy landscape and residue interaction network. All datasets were filtered to include only complexes deposited in the Protein Data Bank after 2020, ensuring they post-date model training sets, and any complex with >30% sequence identity to training data was excluded (verified via PDB70 and MMseqs2 clustering).^62^ For allosteric datasets, an additional pocket-level similarity filter was applied: allosteric pockets with >40% residue identity to a pocket in the training set were excluded.

### Orthosteric Datasets

#### MDT Mainset

The Major Drug Targets (MDT) test set^29^ served as the primary orthosteric reference, comprising 412 complexes from kinases, G protein-coupled receptors, nuclear receptors, and ion channels. Structural challenge was enforced by requiring AlphaFold-predicted apo-pocket deviation from the experimental holo conformation to be >2.0 Å RMSD or pocket LDDT < 0.80, ensuring genuine conformational adaptation. Ligands were restricted to drug-like molecules (200–650 Da), excluding fragments and macrocycles, with no more than ten complexes from any single study to prevent overrepresentation.

#### PLOC Subset

To expand statistical power, we incorporated the Protein–Ligand Orthosteric Competitive (PLOC) subset from a comprehensive dataset of protein interactions and ligand binding pockets.^63^ The PLOC subset contains 1,863 structures with 2,325 binding pockets and 1,647 unique ligands binding to primary, evolutionarily conserved orthosteric sites across 76 Pfam families. Resolution was restricted to ≤3.0 Å. The PLOC complexes were curated to include only competitive ligands occupying the canonical, catalytically relevant binding site. The MDT and PLOC datasets were combined to create a comprehensive orthosteric reference totaling 2,275 complexes.

### Allosteric Datasets

#### Kinase-Specific Allosteric Dataset

Kinase-specific allosteric complexes were derived from the Kinase Conformation Resource (KinCoRe)^64,65^ a geometry-driven classification system enabling unambiguous structural categorization based on DFG motif and αC-helix conformation. Type III (proximal allosteric) inhibitors bind adjacent to the ATP site without entering the cleft (distance >6.0 Å from hinge region, minimum three contacts with back-pocket residues). Type IV (distal allosteric) inhibitors bind >6.5 Å from both the hinge region and the conserved αC-helix Glu(+4) residue. From this resource, we extracted 217 Type III/IV complexes, augmented with a curated X-ray crystallography repository^66^ containing 136 exclusive allosteric kinase inhibitors (resolution ≤3.5 Å, MW ≥250 Da, no primary site occupancy), yielding 353 complexes. Quality-control checks included: (1) resolution ≤3.0 Å, (2) exclusion of ligand atoms with occupancy <0.5, (3) use of only the highest-occupancy conformer for alternate conformations, and (4) manual inspection of a random subset of 50 complexes.

#### Protein–Ligand Allosteric (PLA) Subset

To capture allosteric diversity beyond kinases, we incorporated the Protein–Ligand Allosteric (PLA) subset^63^ containing 1,613 structures with 1,830 binding pockets and 1,277 unique ligands spanning 83 Pfam families across >500 organisms. Resolution was restricted to ≤3.2 Å. All complexes were experimentally validated as authentic allosteric binders with sites spatially distinct from orthosteric pockets. Using UniProt IDs, pocket residue composition, and geometric clustering (Cα RMSD ≤2.0 Å), we identified 182 unique allosteric pockets spanning 115 distinct UniProt entries; for each unique pocket, the highest-resolution representative structure was selected. The kinase-specific allosteric dataset (353 complexes) and the PLA subset (1,613 complexes) were combined to create a comprehensive allosteric ensemble totaling 1,966 complexes. After applying the temporal (post-2020), sequence-similarity (>30% identity excluded), and pocket-level similarity (>40% residue identity excluded) filters, 1,966 initial complexes were filtered down to 503 post-2020, non-redundant complexes. The non-redundant PLA set (182 unique pockets) was retained as a separate, redundancy-controlled benchmark.

### MISATO Dataset

To test whether the frustration–predictability relationship generalizes beyond curated orthosteric and allosteric benchmarks, we incorporated MISATO as a general-purpose benchmark. MISATO is a large machine-learning dataset of protein–ligand complexes derived from PDBbind; it provides experimental binding poses, semiempirical quantum-mechanical refinements of protein and ligand protonation states, and molecular dynamics and electronic-property data for a subset of complexes. The full release contains 19,443 protein–ligand complexes, of which 16,972 have MD simulations. We downloaded the QM.hdf5 and MD.hdf5 files and retained 19,413 ligand IDs from QM.hdf5 with valid 3D SDF representations (19,392 ligands). Complexes that could not be parsed by RDKit were removed. For the docking benchmarks, DiffDock and EquiBind were provided the crystal protein structure and one ligand conformer built from the MISATO QM coordinates. DiffDock was run on 14,029 input pairs; 8,419 produced poses and 5,610 were skipped because RDKit could not parse the ligand. EquiBind was run on 19,392 ligand SDFs and generated 8,565 poses. For the final benchmark, we selected complexes meeting four criteria: (1) resolution ≤ 3.0 Å; (2) complete ligand definitions with valid SMILES; (3) no missing residues in the binding-site region; and (4) MD trajectories of at least 8 ns. Because MISATO was not curated for binding-site type, it may contain ligands at secondary, cryptic, or allosteric sites; we therefore treat it as a general-purpose benchmark. For each selected complex, local frustration analysis was performed on apo and holo structures using the same protocol as for the curated datasets, and binding-site frustration profiles were computed from residues within 4 Å of the ligand.

### AI Co-Folding Architectures

Five co-folding architectures were benchmarked head-to-head following protocols adapted from the PoseBusters benchmark.^25^ All models received identical inputs (protein sequences from PDB structures, ligands in SMILES format, and the same MSAs generated by the standard AF3 pipeline), with no additional structural templates unless specified. For each complex, 25 models were generated on NVIDIA A100 40GB GPUs using 10 recycling steps, 200 diffusion sampling steps, 5 seeds, and 5 diffusion samples per seed, with the ligand–protein chain-pair ipTM score used for ranking. AF3 (v3.0.0)^23^ employs a diffusion-based architecture operating on a unified “cloud of atoms,” simultaneously diffusing protein and ligand coordinates into a synchronized holo-state through iterative refinement, with pairwise attention across all atoms capturing interdependencies between protein conformation and ligand placement. Predictions were generated both with templates enabled (default) and without (AF3-NT). Protenix (v0.3.4)^26^ is an open-source implementation of the AF3 architecture enabling high-throughput inference via unified tokenization of polymers and small molecules while maintaining architectural fidelity to the original diffusion-based framework. Boltz-2 (v2.0.3)^27^ extends diffusion-based co-folding with prediction heads for affinity estimation and binder classification, using confidence-guided sampling to prioritize predictions with high internal consistency. Earlier versions (Boltz-1 v0.4.1, Boltz-1x v1.0.0) were also evaluated. Chai-1 (v0.5.1)^28^ fuses PLM embeddings from ESM-2 with paired attention mechanisms, enabling high structural fidelity even when evolutionary co-variation signal is sparse; predictions were made with the ESM embedding option enabled. DynamicBind^29^ employs an SE(3)-equivariant diffusion graph network that treats docking as a morphological process, representing proteins as coarse-grained residue graphs and ligands as heavy-atom subgraphs, and iteratively adjusting receptor structure from apo to holo states. During inference, DynamicBind generates 20 candidate trajectories per target, scored by a composite metric combining LDDT scores and predicted binding affinities.

### Docking Methods

DiffDock^21^ is a diffusion generative model over the non-Euclidean manifold of ligand poses that redefines docking from a search problem to a generative task. Its SE(3)-equivariant architecture ensures physically consistent predictions with respect to 3D rotation and translation. The model takes ligand SMILES and the native holo protein structure as input, generates amino acid embeddings with ESM2, and updates an initial RDKit^67^ ligand conformation through a diffusion process that models the binding pose distribution without prior knowledge of the protein pockets, making it suitable for blind docking. DiffDock was subsequently updated to DiffDock-L^22^ which provides significant improvement in performance and generalization; both versions were evaluated. For each MISATO complex, 10 independent predictions were generated using default diffusion sampling parameters, ranked by the model confidence score, and both top-1 and top-k metrics were computed. EquiBind^30^ is an SE(3)-equivariant geometric deep learning model performing direct-shot prediction of both the receptor binding location (blind docking) and the ligand bound pose and orientation, optionally coupled with a fast fine-tuning model that adjusts ligand torsion angles via closed-form global minima of the von Mises angular distance to the input atomic point cloud. The model was trained on the PDBBind v2020 dataset (19,443 protein– ligand complexes). For each MISATO complex, a single direct-shot prediction was generated from the protein structure and the ligand molecular graph; both the base EquiBind model and the fine-tuned variant (EquiBind+Q) were evaluated.

### Evaluation Metrics and Alignment Pipeline

Each AI co-folding model employed distinct conventions for file organization, ligand representation, and atom naming. Many ligands possess symmetric substructures or tautomeric states that create multiple geometrically equivalent but atomically distinct binding modes. These model-specific idiosyncrasies necessitated a specialized, adaptive evaluation pipeline. All predicted protein structures were first superimposed onto their experimental reference counterparts using TM-align^68^ for sequence-independent structural alignment. After global alignment, ligands were extracted and evaluated using the OpenStructure toolkit (OST)^69^ which provides direct control over structure alignment, ligand extraction, and RMSD calculation within a unified framework. Ligand Pose RMSD was computed using OST Symmetry-Corrected RMSD Scorer (SCRMSDScorer), which enumerates all possible mappings of model chains onto chemically equivalent target counterparts, calculates RMSD for each symmetry-equivalent atom assignment, and returns the lowest value. LDDT-PLI (Local Distance Difference Test for Ligand–Protein interfaces) evaluated preservation of local atomic distances within the binding pocket, with particular sensitivity to side-chain interactions that backbone RMSD alone cannot capture. Scores range from 0 to 1, with values >0.8 indicating high-fidelity reproduction of interaction geometry.

QS-score (Quality of Structure score) quantified recovery of native contact topology independent of absolute pose geometry, measuring the fraction of native ligand–protein contacts (4.5 Å cutoff) correctly recovered. A QS-score >0.7 generally indicates preservation of the essential binding interaction pattern. Ligand-centric confidence metrics were derived from model-specific outputs. For AF3 and Chai-1, we computed L-pLDDT (arithmetic mean of pLDDT over ligand atoms), L-PAE (average of all PAE elements involving ligand atoms), and mPAE (minimum PAE among elements containing ligand atoms). For Boltz-2, we used ipTM scores and predicted binding affinity (pIC50). Confidence–error correlation was assessed by mapping model-specific confidence scores against actual RMSD.

### Local Frustration Analysis

Local frustration analysis^48–54^ quantified energetic optimality of native interactions by comparing them to decoy ensembles. Two complementary indices were computed for every residue pair in both apo and holo states. Conformational frustration assesses sensitivity to local structural perturbations: for each contact, 1,000 decoy configurations are generated by randomizing residue identities and interatomic distances within native contact geometry, and the native energy is compared to the decoy distribution. Mutational frustration evaluates evolutionary optimality by comparing native interaction energy to 1,000 computational mutations at the same positions. Both frustration indices were expressed as Z-scores:

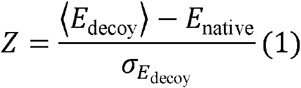

Following validated thresholds,^48–50^ contacts were classified as minimally frustrated (Z > 0.78), highly frustrated (Z < −1.0), or neutrally frustrated (−1.0 ≤ Z ≤ 0.78). Residue-level profiles were generated by computing local density of frustrated contacts within a 5 Å radius; a residue was assigned a dominant frustration category if >50% of its interacting contacts belonged to the same class. Analysis was performed at two levels: across entire complexes for global landscape architecture, and within binding-site regions (residues within 4 Å of ligand). In addition to the standard Frustratometer implementation, frustration calculations were also performed using FrustraMPNN,^59^ a machine-learning-based approach for predicting frustration directly from sequence and structure. This complementary implementation enabled cross-validation of frustration predictions and extension to structural ensembles.

To translate residue-level frustration profiles into quantities comparable across datasets and directly correlated with prediction accuracy, we derived two summary metrics from the binding-site and global-fold frustration fractions tabulated for each dataset and ligand-binding state. Frustration quenching magnitude (ΔMF) quantifies how much the fraction of minimally frustrated contacts at the binding site increases when the ligand binds:

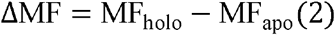

where *MF*_(*apo*)_ and *MF*_(*holo*)_ are the fractions of minimally frustrated residues within the binding-site region in the apo and holo states, respectively. A large positive *ΔMF* indicates that ligand binding resolves energetic conflicts at the binding site and converts a strained region into a minimally frustrated one, generating a steep energetic funnel that enforces geometric convergence. A near-zero *ΔMF* indicates that the binding site is either already in its preferred energetic state in the apo structure or that ligand binding does not resolve local conflicts—in either case, no directional signal is generated. Apo highly frustrated enrichment (*ΔHFapo*) quantifies the extent to which the binding site is a local frustration hotspot relative to the rest of the protein in the ligand-free state:

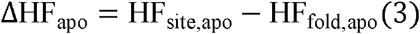

where *HFsite,apo* is the fraction of highly frustrated residues within the binding-site region in the apo state and *HF*_*fold,apo*_ *t*he fraction of highly frustrated residues across the entire protein fold in the same state. A large positive value indicates that the binding site is an energetically conflicted region embedded within an otherwise well-optimized fold—a local frustration hotspot intrinsically poised for reorganization. A negative value indicates that the binding site is less frustrated than the protein average and does not present a distinctive energetic anomaly for detection. Together, the two metrics describe the initial state of the binding site *ΔHF*_apo_ and how that state responds to ligand binding (*Δ MF*).

### Protein Structure Network Analysis

For each benchmark complex, protein structure networks (PSNs) were constructed from the same apo and holo structures used in the frustration analysis. Nodes represent individual residues, with Cα atoms as reference points. Edges were added between residues i and j if the Cα–Cα distance was ≤5.0 Å, matching the contact definition used in the frustration analysis. This residue-level representation of protein structure as a network follows established PSN methodology, in which nodes correspond to residues and edges to non-covalent contacts, and which has been used extensively to relate network topology to protein stability, allosteric communication, and interaction energetics.□□□□□ Each edge was annotated with its conformational frustration Z-score, *zij*, and classified as minimally frustrated (MF; *zij* > 0.78), neutrally frustrated (NF; −1.0 ≤ *zij* ≤ 0.78), or highly frustrated (HF; *zij* < −1.0). For frustration-weighted shortest-path calculations, edge lengths were defined as *dij* = 1.0 if MF, 2.0 if NF, and 10.0 if HF penalizing paths through highly frustrated contacts so that shortest paths preferentially traverse minimally frustrated regions. Network construction and centrality analysis were performed using the RING tools for residue interaction network generation and analysis□^2^□□□ and the resulting networks were further processed with custom Python scripts.

For each residue v, the shortest-path betweenness centrality (SBC) was computed as:

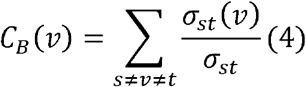

where *σ*_*st*_ is the total number of shortest paths from node s to node t, and *σ*_*st*_*(v)*is the number of those paths passing through v. For an undirected network of N nodes, the normalized value is:

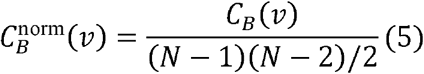

We computed topological SBC with unit edge lengths, and frustration-weighted SBC using the edge lengths defined above. The frustration-weighted SBC was used for all primary analyses reported here.

Z-Score Relative Centrality Analysis (Z-RCA) quantifies whether a residue centrality is statistically exceptional relative to a null model. For each node i, we computed the observed centrality, then generated R = 1,000 degree-preserving random graphs using the configuration model, removed self-loops and multi-edges, and computed the centrality for each random graph r. The Z-RCA score is:

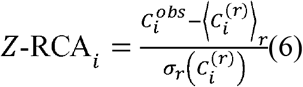

### Binding-Site Aggregation

For each complex, binding-site residues were defined as all residues with at least one atom within 4 Å of the ligand in the holo structure, matching the definition used in the frustration analysis. We then computed:

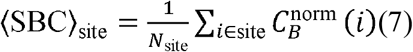

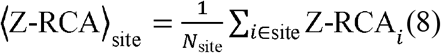

Spearman correlations were calculated between ΔMF and each network metric across all complexes. Orthosteric and allosteric sites were stratified by ΔMF bins (ΔMF > 10%, 5% ≤ ΔMF ≤ 10%, ΔMF < 5%) to test whether network centrality tracks frustration quenching. Logistic regression models using SBC, Z-RCA, and ΔMF as predictors were fit to predict AI prediction success (defined as AF3 oracle ligand RMSD < 2 Å), with 5-fold cross-validation and leave-one-pocket-out validation on the non-redundant PLA set.

### Statistical Analysis

For each model, the best prediction per complex (lowest ligand pose RMSD) was selected from generated models. Average RMSD values represent the mean of best-run RMSDs across all systems in each dataset. Statistical significance of differences between orthosteric and allosteric predictions was assessed using paired two-tailed t-tests (p < 0.05) with Benjamini–Hochberg FDR correction.^75^ Residue-level frustration densities were compared using Mann–Whitney U test, with effect sizes quantified by Cliff’s delta.^76^ Spearman’s rank correlation was computed between per-residue frustration Z-scores and per-residue model confidence scores. Model performance was compared across the following dimensions: (1) co-folding models vs. docking models (DiffDock, EquiBind), (2) orthosteric vs. allosteric complexes, and (3) complexes stratified by binding pocket depth, ligand properties, and MD-derived flexibility metrics. Confidence–error correlations were computed for DiffDock predictions to assess the reliability of the model confidence estimates. The full evaluation pipeline, including all scripts for input preparation, model execution, output postprocessing, and statistical analysis, is available in our open-source repository with detailed documentation for community adoption and extension.

## Results

### Binding-Site Predictability as a Property of Energetic Landscape Organization

AI co-folding and docking models predict ligand binding poses with high accuracy at classical orthosteric sites but systematically lose accuracy at allosteric and cryptic sites. This gap has been reproduced across major architectures and metrics.^47^ Whether orthosteric and allosteric sites represent discrete classes of prediction problem or whether they differ in a physical property that varies across binding sites remains an open question. To address these questions, we established an integrated analytical pipeline (Figure 1).

**Figure 1.**
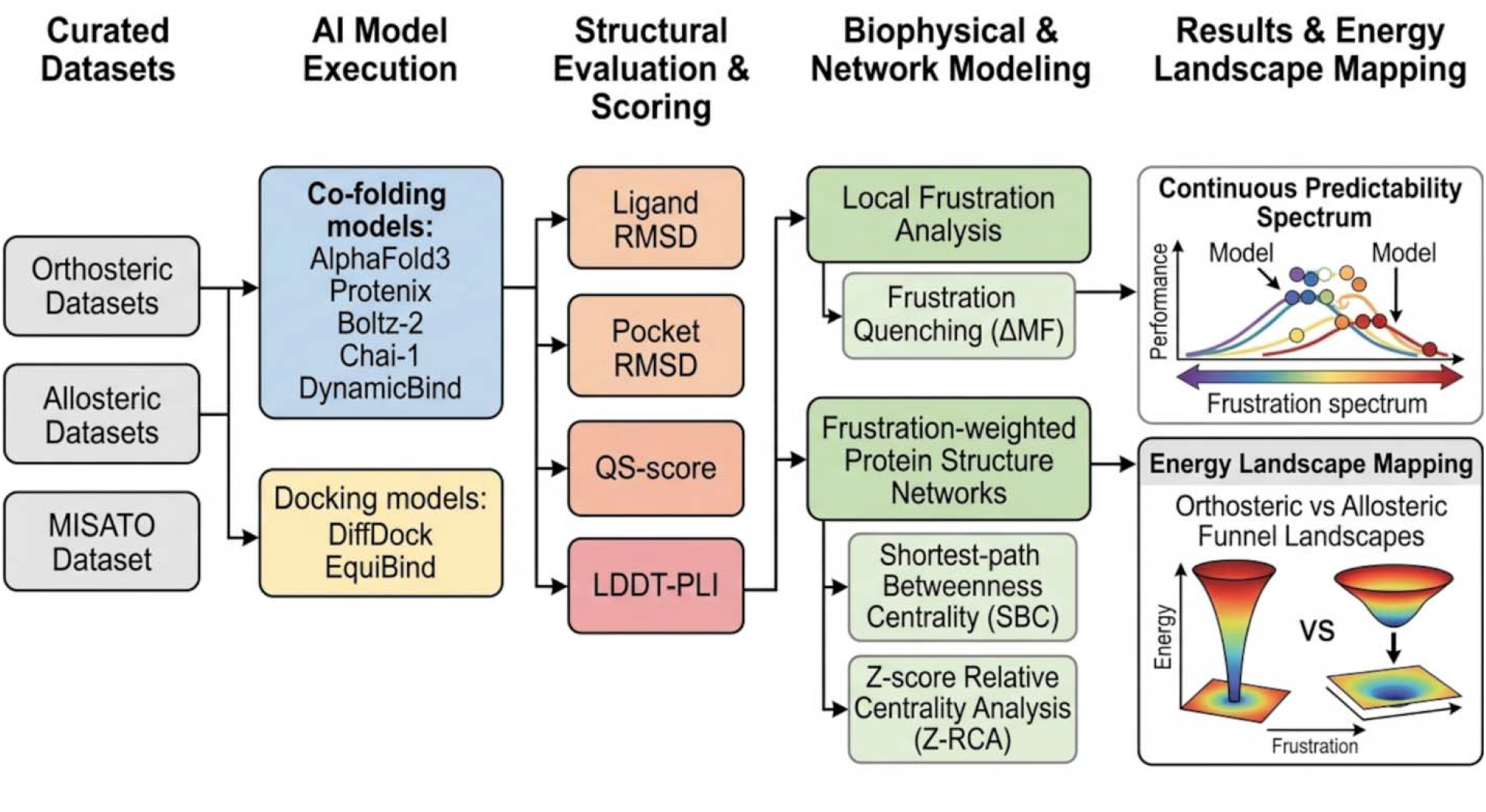
Overview of the Benchmarking, Evaluation, and Analysis Pipeline. Schematic workflow of the study. Left (Datasets): the orthosteric and allosteric datasets are assembled and curated, together with the general-purpose MISATO benchmark. Center (Pipeline): inputs are processed into standardized formats, the seven co-folding and docking models are executed on identical inputs, raw outputs are converted into a uniform representation, predictions are scored against experimental references, scores are aggregated across complexes and stratified by ΔMF and network centrality, and figures and tables are generated. Top right (Models): the pipeline forks into the evaluated architectures. Bottom right (Evaluation Metrics): each predicted structure is scored against the experimental reference using complementary metrics.

First, across four complementary metrics, we confirm that the orthosteric–allosteric accuracy gap is robust against benchmark confounders and transcends the co-folding paradigm. Next, extending this framework to the broad MISATO benchmark—covering diverse interfaces and MD-derived cryptic sites—reveals local frustration quenching as the primary physical driver of prediction accuracy. Finally, we show how binding landscape topology and site communication architecture organize AI model performance along two fundamental physical axes.

### The Orthosteric–Allosteric Gap and Its Structural Signature

We first establish the orthosteric–allosteric performance gap^47^ and its structural signature across the four complementary metrics that organize the rest of this work: ligand pose RMSD, QS-score, LDDT-PLI, and success rate at 2.0 Å (Table 1). Across the five co-folding models, ligand pose accuracy sorts into two regimes. Orthosteric predictions converge on native geometries, with mean ligand RMSD 2.3–2.5 Å for AF3, Protenix, and Chai-1 and 3.8–4.1 Å for Boltz-2 and DynamicBind. Allosteric predictions achieve mean ligand RMSD = 5.2–6.8 Å for every model.

**Table 1.** Performance metrics of deep learning docking models and joint biomolecular prediction frameworks, stratified by orthosteric and allosteric binding sites.

| **Model** | **Mean Ligand RMSD** | **QS-score** | **LDDT-PLI** | **Success @ 2 Å** |
| --- | --- | --- | --- | --- |
|  | **(Å) Ortho / Allo** | **Ortho / Allo** | **Ortho / Allo** | **Ortho / Allo** |
| AF3 | 2.3 / 6.1 | 0.94 / 0.72 | 0.95 / 0.74 | 82% / 29% |
| Protenix | 2.5 / 6.2 | 0.94 / 0.70 | 0.94 / 0.72 | 81% / 26% |
| Chai-1 | 2.4 / 5.2 | 0.93 / 0.73 | 0.95 / 0.73 | 80% / 31% |
| Boltz-2 | 3.8 / 6.8 | 0.91 / 0.68 | 0.88 / 0.71 | 74% / 23% |
| DynamicBind | 4.1 / 5.2 | 0.90 / 0.75 | 0.80 / 0.75 | 72% / 33% |

\* Mean Ligand RMSD: Average heavy-atom RMSD across all target predictions in the respective site category. Success Rate @ 2.0 Å is defined as the percentage of top-ranked predicted ligand poses achieving a heavy-atom RMSD $\leq 2.0\text{\AA}$ relative to the experimental crystal structure.

The gap is significant in both mean RMSD and in the fraction of predictions reaching near-native accuracy as the success prediction rates at RMSD = 2.0 Å fall from 72–82% on orthosteric sites to 23–33% on allosteric sites. Notably, the ranking and performance of AI co-folding models on orthosteric sites do not necessarily track their ranking on allosteric sites. In fact, AF3 leads on orthosteric sites but drops behind Chai-1 and DynamicBind on allosteric sites while DynamicBind performs last on orthosteric sites but matches Chai-1 on allosteric sites.

Table 1 exposes the structural signature of the gap. QS-score, which reports on native contact recovery independently of absolute pose geometry, exceeds 0.90 on orthosteric complexes and remains moderately high at 0.70–0.85 on allosteric complexes despite ligand RMSD exceeding 6 Å. The two metrics are decoupled at allosteric sites: a model recovers the correct contact set while placing the ligand several angstroms from its native position. Models recover the identity of the contacting residues but not the geometry of their arrangement. LDDT-PLI refines this decoupling into three tiers by probing the intermediate scale, local interaction fidelity including side-chain geometry. Orthosteric LDDT-PLI is tightly distributed above 0.90, with medians >0.95 for AF3, Chai-1, and Protenix. Allosteric LDDT-PLI broadens downward, with medians of 0.70–0.80 and long tails below 0.5. The three-tier pattern is: contact identity is largely preserved (QS-score 0.70–0.85), local geometric arrangement is partially preserved (LDDT-PLI 0.60–0.80), and global geometric arrangement is lost (ligand RMSD >6 Å). LDDT-PLI reports on the specific spatial scale at which allosteric prediction fails. The gap and its structural signature are robust to the ways the benchmarks could be confounded. They persist across the non-redundant PLA set under both oracle and average selection strategies and independently on MISATO (Tables S1, S2). The mean orthosteric ligand RMSD never exceeds 3.0 Å and the mean allosteric ligand RMSD never falls below 5.2 Å suggesting that the gap is a property of the binding-site classes themselves.

### Two Docking Models Occupy Opposite Extremes of the Sampling Spectrum

The five initial AI models are co-folding architectures that jointly predict protein and ligand coordinates, so the gap may reflect a limitation of the co-folding paradigm rather than of the binding sites. Second, the curated benchmarks of orthosteric and allosteric complexes may not reveal how prediction accuracy varies across the broader range of binding sites. To address this, we compared two docking models that occupy opposite extremes of the sampling spectrum on a shared control set (MISATO dataset). DiffDock, a generative diffusion model that samples and ranks multiple candidate poses, and EquiBind, a rigid regression model that produces a single pose in one forward pass. DiffDock generates ten independent predictions per complex and ranks them by its confidence score. EquiBind generates one. If the gap were a property of co-folding, neither model should reproduce it. If the gap depended on the capacity to generate and rank multiple poses, DiffDock should reproduce it and EquiBind should not. The distributions in Figure 2 support the second interpretation. DiffDock produces a narrow pose RMSD distribution centered near 0–1 Å, with an interquartile range of approximately 1.5 Å and a short upper tail. EquiBind produces a broad distribution centered at 3–5 Å with a tail extending beyond 10 Å (Figure 2A). The two distributions differ in shape as well as in central tendency: DiffDock is unimodal and tightly peaked, whereas Equibind is broad and heavy-tailed, with substantial weight in the 5–10 Å range that DiffDock does not occupy.

**Figure 2.**
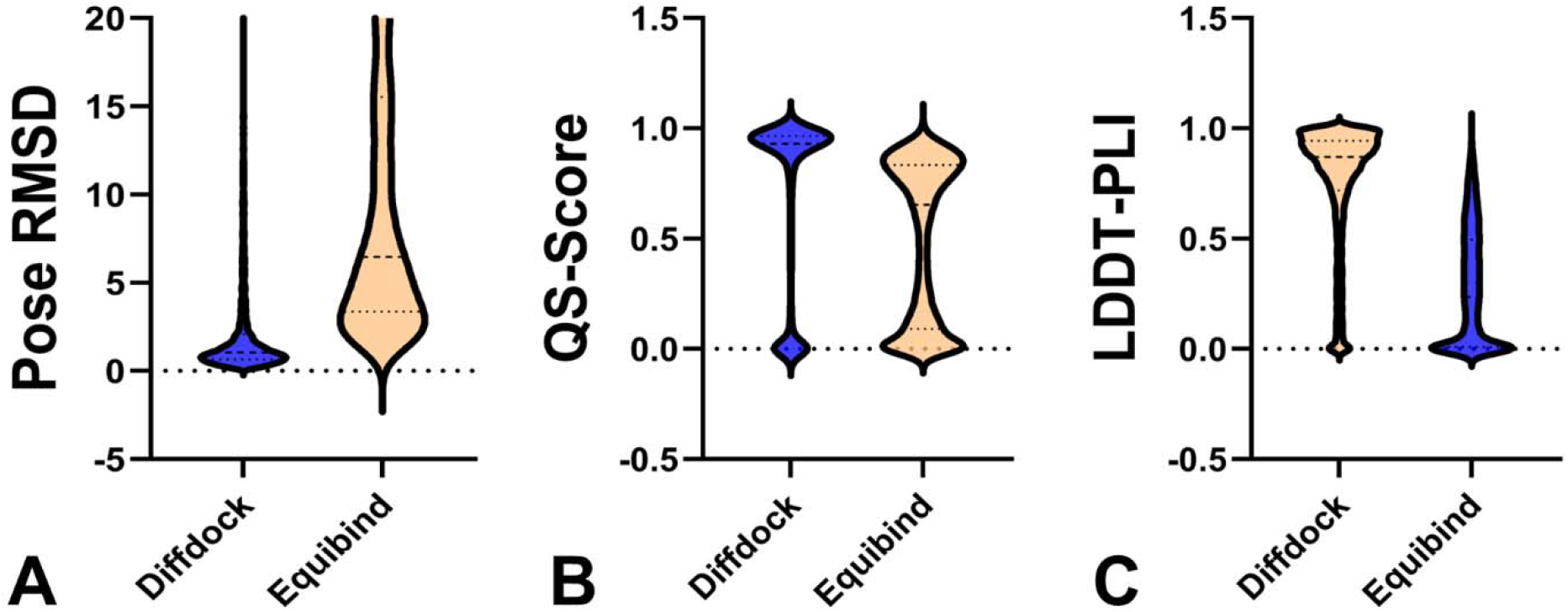
Comparison of Two Docking Models at Opposite Extremes of the Sampling Spectrum on the MISATO Set. Violin plots comparing DiffDock (blue) and EquiBind (orange) across three complementary metrics : pose RMSD (A), QS-score (B), and LDDT-PLI (C). Each violin shows the full distribution of per-complex values; dotted lines inside each violin indicate the median and quartiles. DiffDock distributions are concentrated at the high-accuracy end, with narrow interquartile ranges and short lower tails. Equibind distributions are broad and shifted toward lower accuracy, with substantial weight in the poorly performing range.

The QS-score distributions follow the same pattern (Figure 2B). DiffDock has a median above 0.95 with a tight distribution and a short lower tail. EquiBind has a median near 0.85 with a lower tail extending toward 0.0. LDDT-PLI distributions show the same contrast (Figure 2C). DiffDock is tightly distributed near 1.0 with negligible spread below 0.5; EquiBind spans the full 0–1 range with a broad peak near 0.8 and a long lower tail. The gaps between the two models are not equal across metrics. The gap is largest for pose RMSD (approximately 4 Å in median), smaller for QS-score (approximately 0.10 in median), and smallest for LDDT-PLI (approximately 0.15 in median). This ordering indicates that the two models differ most in their capacity to reach the correct global geometry and least in their capacity to reproduce local interaction geometry once a pose has been generated. Two mechanisms are consistent with this ordering. Under the first, sampling capacity sets the accuracy ceiling for global geometry, while local interaction fidelity is limited by the model representation of the binding pocket. Under the second, both metrics are limited by sampling capacity, but global geometry is more sensitive to the number of candidate poses than local geometry. The analysis suggests that the distinction between the two docking models is not the difference between co-folding and docking. DiffDock is a docking model and yet matches the co-folding performance profile. The relevant distinction is between a model that generates and ranks multiple candidate poses and one that does not. Symmetry-corrected RMSD tools such as LigRMSD,^77^ DockRMSD,^78^ and Spyrmsd^79^ were used to confirm that the contrast between the two models is not an artifact of any single RMSD implementation. All three tools produce the same ordering on the MISATO set, with the same separation between the DiffDock and EquiBind distributions across pose RMSD, QS-score, and LDDT-PLI.

### A General-Purpose Benchmark Reveals the Full Range of Prediction Accuracy

The benchmarks used in the preceding sections were curated for site type. The MDT and PLOC sets were assembled to represent canonical orthosteric pockets. The KinCoRe and PLA sets were assembled to represent authentic allosteric binders that are spatially distinct from the orthosteric site. This curation produces the pronounced gap in Table 1, but it also restricts the benchmark to two specific site categories. It does not tell us how prediction accuracy is distributed across the broader range of protein–ligand interfaces that occur in the structural record.

MISATO is a broad structural-biology dataset derived from PDBbind that provides experimental protein–ligand complexes together with semiempirical quantum-mechanical refinements of protein and ligand protonation states, and molecular dynamics trajectories for a substantial subset of the entries.^61^ It contains the full spectrum of protein–ligand interfaces that occur in the structural record. These include canonical orthosteric pockets, distal allosteric pockets, cofactor-binding sites, and unclassified or multi-functional interfaces. The MD data extend this spectrum further. Each trajectory samples conformational fluctuations and exposes transient and cryptic pockets that may not be visible in the static crystal structure. Cryptic sites are of particular interest here because their absence in the deposited conformation means that a model must infer the pocket from the sequence and the surrounding fold rather than read it off the input coordinates. MISATO therefore allows the analysis to sample binding sites across the full range of conformational accessibility, from sites that are constitutively present to sites that appear only in the ensemble. Figure 3 shows the distribution of predicted pose RMSDs for each of the seven models on the MISATO benchmark.

**Figure 3.**
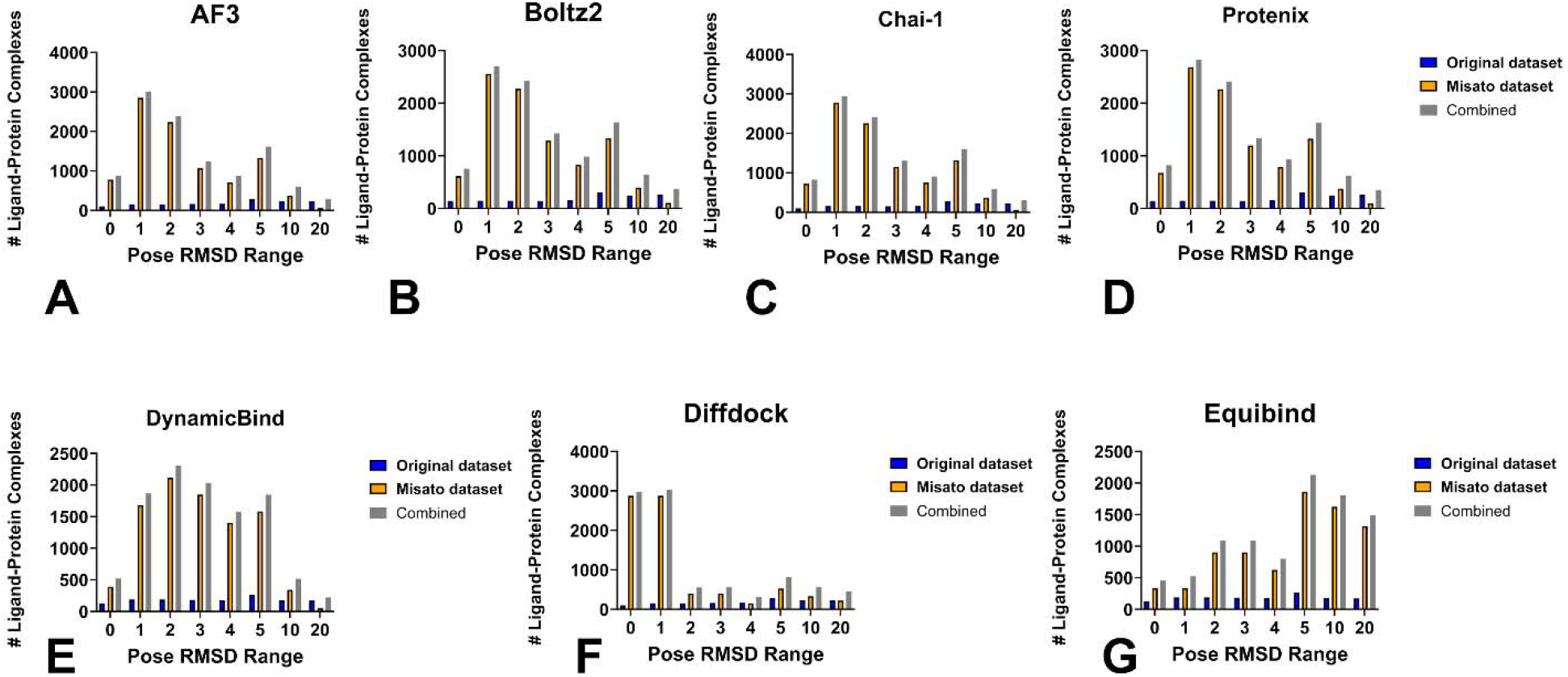
Ligand Pose RMSD Distributions on the MISATO Benchmark. Bar charts showing the number of ligand–protein complexes in each pose RMSD bin (0, 1, 2, 3, 4, 5, 10, and 20 Å) for each of the seven models evaluated on MISATO. Panel A, AF3; Panel B, Boltz-2; Panel C, Chai-1; Panel D, Protenix; Panel E, DynamicBind; Panel F, DiffDock; Panel G, EquiBind. The four top co-folding models, DynamicBind, and DiffDock produce distributions with a dominant mode at 1–3 Å and a monotonic decay toward higher RMSD. The distributions span the full range from sub-angstrom to more than 20 Å and are similar in shape across architectures. EquiBind produces a distribution that peaks at 5–10 Å and does not produce a low-RMSD mode; the origin of this difference is analyzed in the section that follows.

Three features of the MISATO distributions are relevant to the analysis that follows. First, for every model that samples and ranks multiple candidate poses, the distribution of predicted RMSDs spans a wide range, from sub-angstrom accuracy at the low end to more than 10 Å at the high end, with a peak at low RMSD and a long right tail. The fraction of complexes predicted at near-native accuracy varies from 29% for DiffDock to 57% for AF3. Second, the shape of the distribution is similar across the six sampling-capable models. AF3, Boltz-2, Chai-1, Protenix, and DiffDock each show a peak at 1–2 Å with monotonic decay (Figure 3A–D and 3F), and DynamicBind shows the same shape with a slightly broader peak at 2–3 Å (Figure 3E). The similarity of the shapes across architectures that differ in training data, objective, and co-folding machinery indicates that the variation in prediction accuracy is not specific to any one model. Third, EquiBind departs from this pattern (Figure 3G). Its distribution peaks at 5–10 Å, contains substantial weight at 10 and 20 Å, and does not produce a low-RMSD mode.

Table 2 demonstrates that ligand pose prediction accuracy on the general-purpose MISATO benchmark is primarily governed by a model sampling capacity and its ability to navigate the local energy landscape, rather than the co-folding paradigm alone. The four high-capacity co-folding architectures (AF3, Chai-1, Protenix, and Boltz-2), which each sample 25 candidate poses per complex, form a tightly converged top tier: they achieve success rates at 2.0 Å of 53%–57%, RMSD peaks within 1–2 Å, high contact topology preservation (mean QS-score = 0.84–0.86), and strong local interaction fidelity (median LDDT-PLI > 0.88 to > 0.90). Crucially, DiffDock demonstrates that co-folding is not strictly required to achieve high local interaction accuracy; as a generative docking model generating 10 candidate poses, DiffDock achieves a tight RMSD peak of 0–2 Å and a median LDDT-PLI > 0.90 that matches the top co-folding models, though its overall success rate (29%) and contact recovery (QS-score = 0.71) are constrained by the lack of receptor coordinate co-adaptation. DynamicBind occupies an intermediate position (41% success rate @ 2.0 Å, RMSD peak at 2–3 Å, QS-score = 0.75) where its partial induced-fit flexibility offers receptor adaptation at the expense of top-end geometric precision on steep funnels. In sharp contrast, EquiBind’s single shot regression approach (sampling capacity = 1) fails across all metrics, producing an RMSD peak at 5–10 Å, a mean QS-score of 0.38, and a success rate of just 8%.

**Table 2.**
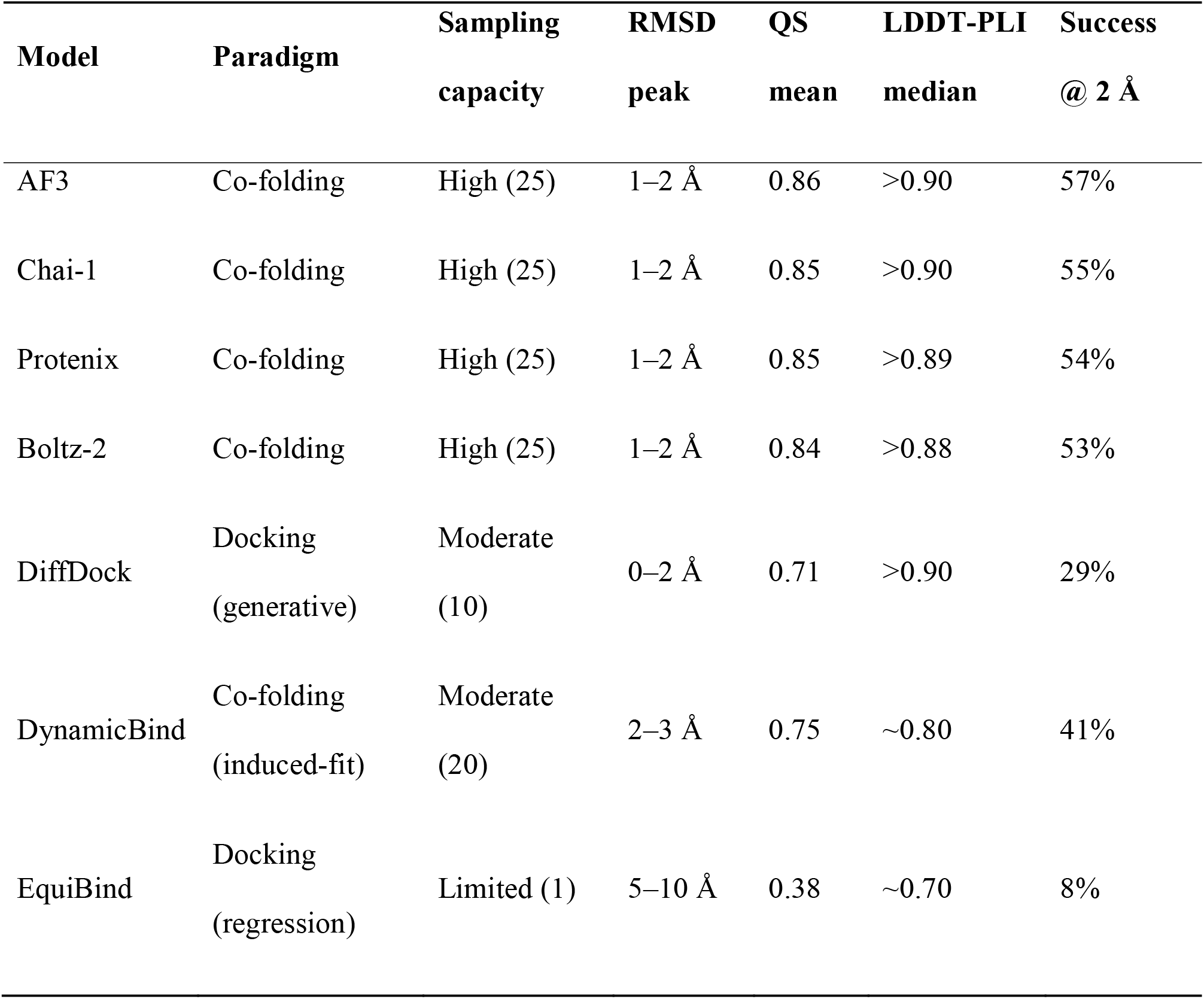
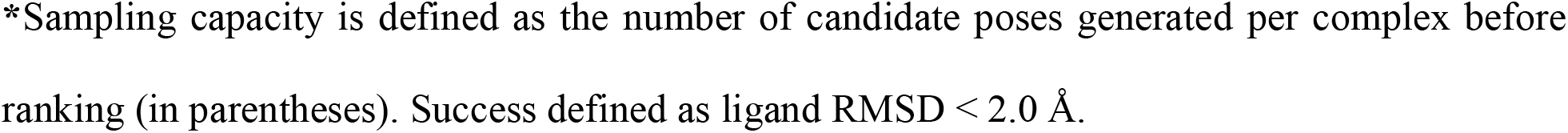
Performance of All Seven Models on the MISATO Benchmark.^*^

| **Model** | **Paradigm** | **Sampling**<br>**capacity** | **RMSD**<br>**peak** | **QS**<br>**mean** | **LDDT-PLI**<br>**median** | **Success**<br>**@ 2 Å** |
| --- | --- | --- | --- | --- | --- | --- |
| AF3 | Co-folding | High (25) | 1–2 Å | 0.86 | >0.90 | 57% |
| Chai-1 | Co-folding | High (25) | 1–2 Å | 0.85 | >0.90 | 55% |
| Protenix | Co-folding | High (25) | 1–2 Å | 0.85 | >0.89 | 54% |
| Boltz-2 | Co-folding | High (25) | 1–2 Å | 0.84 | >0.88 | 53% |
| DiffDock | Docking<br>(generative) | Moderate<br>(10) | 0–2 Å | 0.71 | >0.90 | 29% |
| DynamicBind | Co-folding<br>(induced-fit) | Moderate<br>(20) | 2–3 Å | 0.75 | ~0.80 | 41% |
| EquiBind | Docking<br>(regression) | Limited (1) | 5–10 Å | 0.38 | ~0.70 | 8% |

\*Sampling capacity is defined as the number of candidate poses generated per complex before ranking (in parentheses). Success defined as ligand RMSD < 2.0 Å.

Together, these results establish that generating and ranking candidate poses is an essential architectural requirement for navigating energy landscapes; without explicit pose sampling, single-shot models cannot locate global minima or resolve binding geometries across uncurated protein–ligand interfaces.

### Frustration Quenching is a Property of the Binding Landscape Topology

Because the six sampling-capable models produce distributions of similar shape despite large differences in training data and objective, the determinant of where a given complex falls on the accuracy range is more likely to be a property of the binding site than a general capability of the model. The frustration analysis in the next section tests this directly. We propose that what distinguishes prediction accuracy of binding sites and ligand poses is the topology of the local energy landscape at the binding interface. Local frustration analysis provides a residue-level readout of this topology, and two summary metrics translate the readout into quantities that can be compared across datasets and correlated with prediction accuracy. The frustration quenching magnitude *ΔMF* measures how much the fraction of minimally frustrated contacts at the binding site increases when the ligand binds. The apo highly frustrated enrichment *ΔHF apo* measures the density of highly frustrated contacts at the binding site relative to the rest of the protein in the unbound state.

The two metrics describe different aspects of the binding landscape. Δ*HF*_apo_ reports on how well organized the landscape is around the binding site before the ligand arrives. A high positive value indicates that the site is a locally frustrated region embedded in a more ordered surrounding fold, which is the signature of a landscape that is poised to reorganize upon binding. A negative or near-zero value indicates that the site is no better organized than the protein average and presents no distinctive structural feature for the ligand to exploit. *ΔMF* reports on how the landscape reorganizes when the ligand binds. A large positive value indicates that the ligand-induced contact set populates a contiguous minimally frustrated subnetwork spanning the binding pocket, which produces a coherent directional gradient across the interface. A near-zero value indicates that the ligand-bound landscape is no more ordered than the ligand-free landscape, so no directional gradient emerges. When both metrics are large and positive, the binding site corresponds to a well-defined funnel with a single dominant minimum. When both are near zero, the binding site corresponds to a flat landscape with multiple shallow minima.

The conformational and mutational frustration profiles across the three datasets are reported in Tables S3 and S4. Four features of these profiles place the three datasets along a single ordering. First, the global fold distributions are nearly invariant across datasets and across apo and holo states, confirming that the discriminating signal is local to the binding interface. Second, orthosteric binding sites combine a high *ΔHF apo* with a large *ΔMF*. Third, allosteric binding sites combine a negative or near-zero *HF apo* with a near-zero *ΔMF*. Fourth, MISATO binding sites occupy intermediate values of both metrics where *ΔMF* increases monotonically from allosteric through MISATO to orthosteric, and *ΔHF apo* follows the same ordering. To evaluate whether local energetic funneling directly governs docking accuracy, we stratified AF3 oracle predictions across the dataset (*n* = 9,407) into four quartiles according to their frustration quenching magnitude (ΔMF; Table 3). Prediction accuracy decayed monotonically from the highest frustration quenching quartile (Q1, ΔMF *>* 18%) to the lowest (Q4, ΔMF < 6%). Complexes in Q1 achieved a mean ligand RMSD of 2.6 ± 1.0 Å and a 74% success rate (< 2.0 Å RMSD), whereas complexes in Q4 exhibited a mean ligand RMSD of 5.8 ± 1.8 Å with a success rate of only 24%. Interface contact fidelity mirrored this geometric trend, with QS-score dropping from 0.92 to 0.73 and LDDT-PLI decreasing from 0.90 to 0.67 across quartiles. A non-parametric Kruskal–Wallis *H* test confirmed that performance variations across ΔMF quartiles were highly statistically significant across all four metrics (*H* = 1,480−2,150, all *p*< 1 × 10^−100^; Table 3), exhibiting strong effect sizes (*∈*^2^ = 0.16–0.23). All metrics remained significant after Benjamini–Hochberg false discovery rate correction (*q* < 0.001). Furthermore, post-hoc Dunn’s pairwise testing with Bonferroni correction confirmed that differences between every pair of quartiles were statistically significant for ligand RMSD (all *p*< *0*.*05*), demonstrating a robust link between frustration quenching and predictive accuracy. These findings demonstrate that local frustration quenching is a statistically robust determinant of docking pose accuracy and contact fidelity, establishing energetic funneling as a key driver of protein–ligand binding predictability.

**Table 3.**
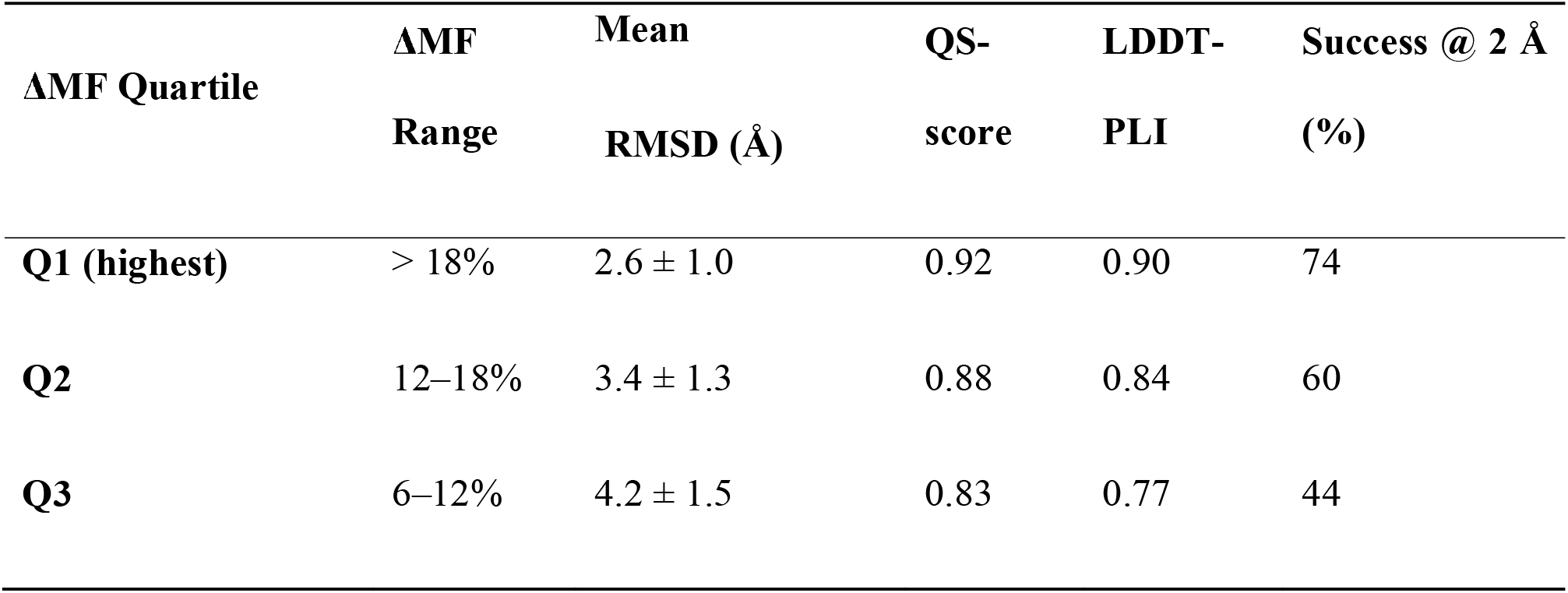
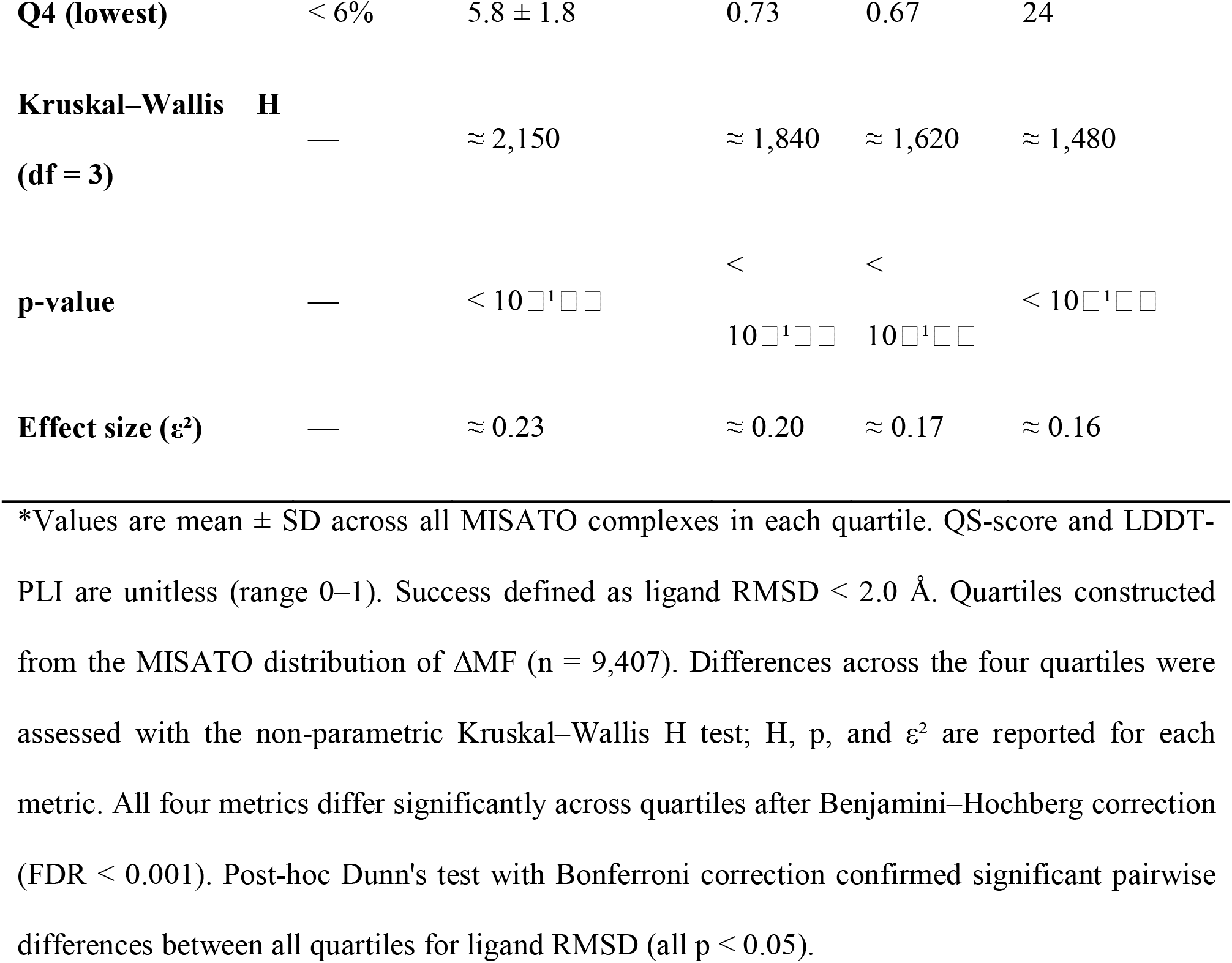
Prediction Accuracy as a Function of Frustration Quenching Magnitude (AF3, Oracle Selection, MISATO).^*^

Figure 4 and Table 4 quantify how effectively different AI architectures convert local energy landscape gradients into geometric pose precision across the MISATO benchmark by introducing Effective ΔMF—a metric that inverts the physical relationship between local frustration quenching (ΔMF) and accuracy to calculate the operating point implied by a model mean ligand RMSD. Table 4 partitions seven evaluated models into distinct performance tiers, showing that high-capacity co-folding architectures (AF3, Chai-1, Protenix, and Boltz-2) tightly cluster in the 9.1%-12.5% Effective ΔMF range (Mean RMSDs of 3.6Å-4.1 Å) by utilizing multi-pose sampling (25 candidate poses) and ranking to navigate steep energy funnels. Conversely, architectures with restricted or absent sampling operate at substantially lower operating points, moving from induced-fit flexible co-folding in DynamicBind (5.8%, 4.8 Å) and rigid generative diffusion in DiffDock (4.4%, 5.2 Å) down to single-shot deterministic regression in EquiBind (1.2%, 7.2Å).

**Table 4.**
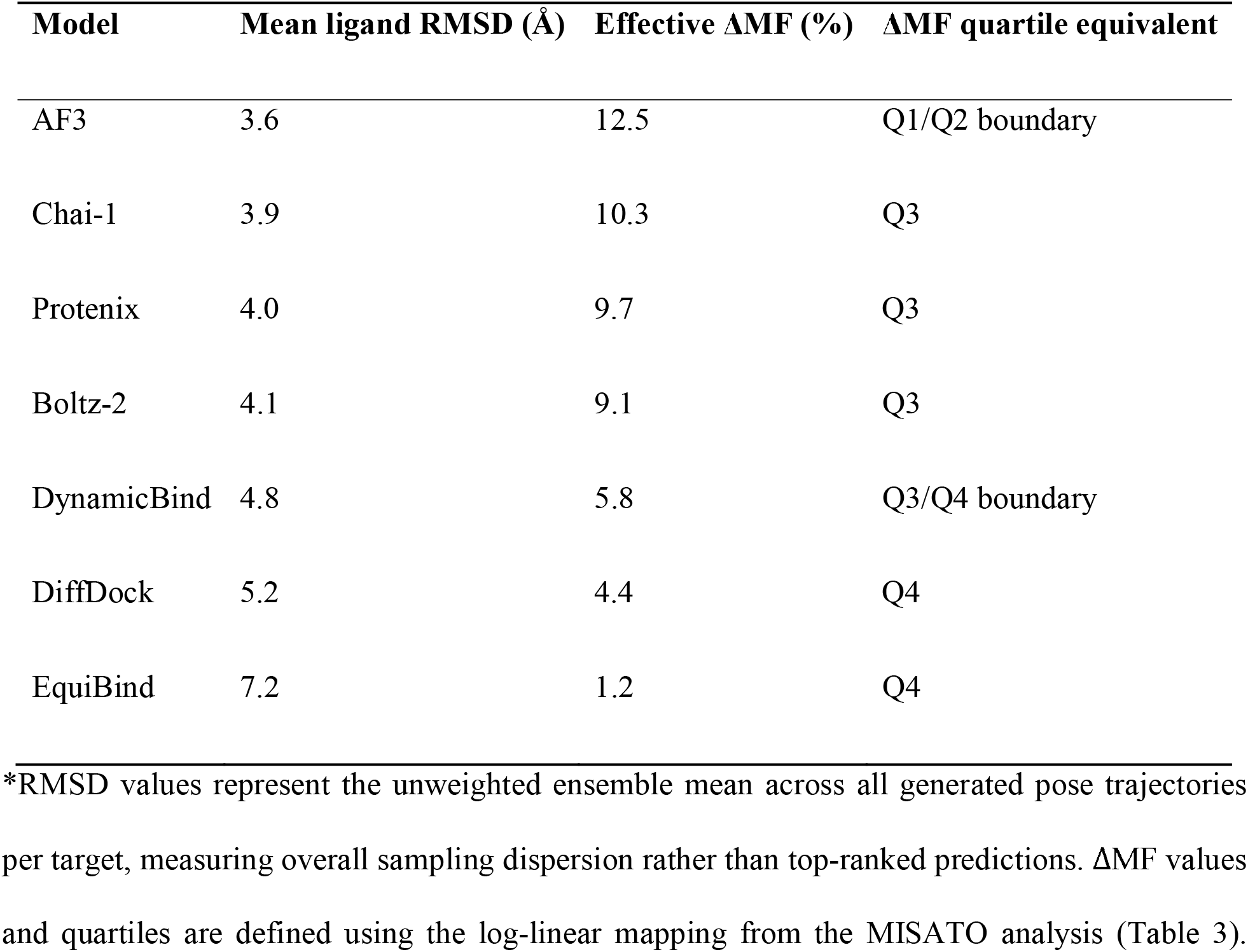
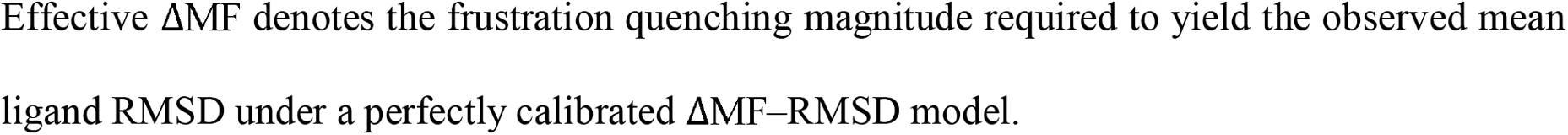
ΔMF Partition Derived from Mean Ligand RMSD for Each Model on MISATO Dataset.

| Model | Mean ligand RMSD (Å) | Effective ΔMF (%) | ΔMF quartile equivalent |
| --- | --- | --- | --- |
| AF3 | 3.6 | 12.5 | Q1/Q2 boundary |
| Chai-1 | 3.9 | 10.3 | Q3 |
| Protenix | 4.0 | 9.7 | Q3 |
| Boltz-2 | 4.1 | 9.1 | Q3 |
| DynamicBind | 4.8 | 5.8 | Q3/Q4 boundary |
| DiffDock | 5.2 | 4.4 | Q4 |
| EquiBind | 7.2 | 1.2 | Q4 |

\*RMSD values represent the unweighted ensemble mean across all generated pose trajectories per target, measuring overall sampling dispersion rather than top-ranked predictions. $\Delta$ MF values and quartiles are defined using the log-linear mapping from the MISATO analysis (Table 3).
Effective $\Delta$ MF denotes the frustration quenching magnitude required to yield the observed mean ligand RMSD under a perfectly calibrated $\Delta$ MF–RMSD model.

**Figure 4.**
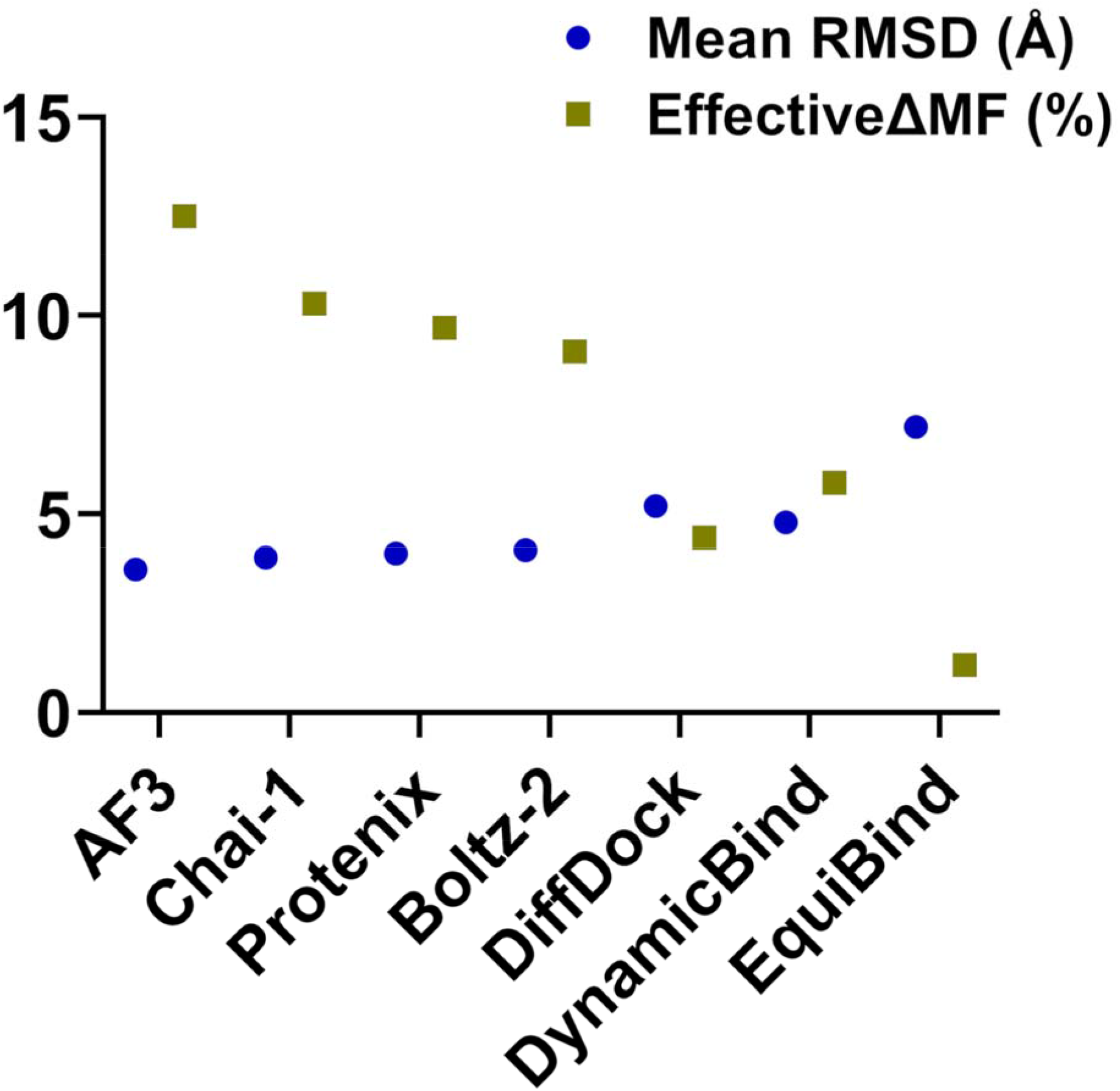
Mean Ligand RMSD and Effective ΔMF for All Seven Models on the MISATO Benchmark. Scatter plot of mean ligand RMSD (blue circles) and effective frustration quenching magnitude ΔMF (olive squares) for each of the seven models evaluated on the MISATO benchmark. Both quantities are read against the y-axis: mean ligand RMSD in Å (0– 15) and effective ΔMF in percent (0–15). Effective ΔMF is not a property of the binding sites but of each model average operating point on the ΔMF–RMSD curve; it is obtained by inverting the log-linear mapping in Table 3 for the model observed mean ligand RMSD.

Figure 4 visually illustrates this strict inverse relationship between pose error and Effective ΔMF, establishing that candidate pose sampling capacity governs funnel accessibility, while receptor coordinate co-adaptation dictates exploitation efficiency once inside a funnel. Single-shot models like EquiBind collapse on flat landscapes because they cannot sample alternative minima, whereas high-capacity co-folding models successfully leverage steep frustration gradients to resolve global minima. Ultimately, this quantitative mapping proves that AI model failures at unquenched, low-ΔMF binding sites (such as allosteric or cryptic pockets) reflect physical limits—specifically, flat local energy landscapes lacking dominant directional funnels—rather than superficial software defects. To examine how each model accuracy is composed across the frustration spectrum, we partitioned the MISATO benchmark into four ΔMF quartiles and evaluated both the complex counts and performance trajectories for all seven architectures (Figure 5 and Table 5). The five co-folding models AF3, Chai-1, Boltz-2, DynamicBind and Protenix exhibit roughly uniform distributions across the quartiles, indicating that their baseline accuracy tracks the underlying benchmark composition and decays smoothly as frustration quenching diminishes (Figure 5). This uniform spread confirms that high-capacity co-folding models do not selectively overperform on specific subsets, but rather experience a generalized, continuous degradation in geometric accuracy as the binding landscape flattens from Q1 (ΔMF *>* 18%) to Q4 (ΔMF < 6%).

**Table 5.** Distribution of Complexes Across ΔMF Quartiles for Each Model.

| **Model** | **Q1 (>18%)** | **Q2 (12–18%)** | **Q3 (6–12%)** | **Q4 (<6%)** |
| --- | --- | --- | --- | --- |
| AF3 | 28% | 26% | 25% | 21% |
| Chai-1 | 25% | 25% | 26% | 24% |
| Protenix | 24% | 25% | 26% | 25% |
| Boltz-2 | 23% | 25% | 26% | 26% |
| DiffDock | 38% | 22% | 20% | 20% |
| DynamicBind | 20% | 23% | 27% | 30% |
| EquiBind | 10% | 12% | 25% | 53% |

\*Percentages are estimated by mapping each model’s ligand RMSD distribution onto the $\Delta MF$ quartile boundaries derived from the log-linear fit. Q1 = highest frustration quenching; Q4 = lowest. The sum of each row is 100%. Values are approximate and intended for comparative rather than absolute interpretation.

**Figure 5.**
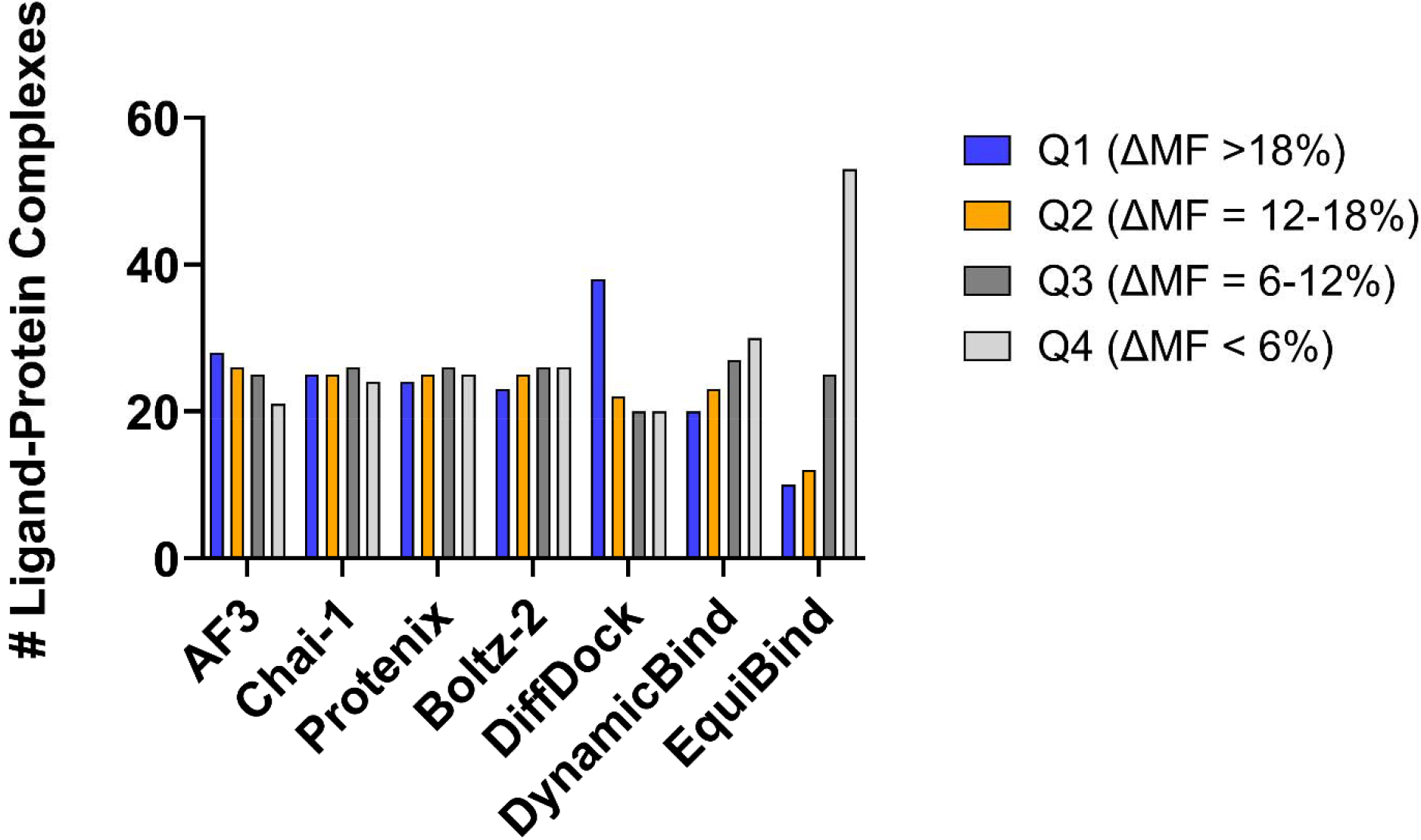
Model performance and complex distribution across local frustration quenching (ΔMF) quartiles. Bar chart displaying the distribution and count of protein–ligand complexes evaluated across AI models—including co-folding architectures (AF3, Protenix, Boltz-2, Chai-1, DynamicBind) and docking methods (DiffDock, EquiBind)—stratified by the magnitude of frustration quenching (ΔMF = MF_holo_ − MF_apo_). Complexes are categorized into four frustration quartiles: Q1 (ΔMF > 18%, strong frustration quenching / well-defined energy funnels), Q2 (ΔMF ≈ 12-18%), Q3 (ΔMF ≈ 6-12%), and Q4 (ΔMF < 6%, minimal frustration quenching / highly frustrated allosteric sites).

Two models depart from the uniform distribution, and their departures reflect different architectural limitations. DiffDock is skewed toward Q1 (highest ΔMF, ≈ 38%), meaning its successful predictions concentrate at the sites where the ligand-induced funnel is steepest. Its sampling capacity allows it to converge accurately on well-defined funnels, but not on flat landscapes. DynamicBind is mildly skewed toward Q4 (lowest ΔMF, ≈ 30%). Its partial induced-fit mechanism allows the receptor to adapt to some ligands even at low-ΔMF sites, so it does not collapse entirely in the flat regime, but the adaptation is not sufficient to fully exploit the steep funnels at high ΔMF. The two skews therefore have different origins: DiffDock is a direct consequence of depending on a funnel that is sometimes absent, while DynamicBind reflects a mechanism that partially compensates for flat landscapes but at the cost of lower performance at the steep end.

The effective ΔMF values, the quartile distributions in Table 5, and the performance trajectories depicted in Figure 5 collectively establish that the seven models can be organized along two orthogonal axes governed by binding landscape topology: funnel accessibility (whether a model can engage with the local gradient) and exploitation efficiency (how precisely it resolves the global minimum once inside the funnel).

### The Two Axes Are Landscape Properties That Organize the Behavior of All Seven Models

We found that model performance across energetic landscapes is governed by two distinct axes: funnel accessibility and exploitation efficiency. In target sites characterized by low frustration quenching ΔMF, the energy landscape constraints severely limit pose sampling across all architectures, with only 8.8% of total predictions reaching a high Quality Score (QS > 0.9). By contrast, EquiBind’s baseline performance across the entire benchmark dataset reflects an overall success rate of 8.0% (RMSD < 2.0 Å) and a mean Quality Score of 0.38 (Table 2). When evaluated specifically within the low-ΔMF subset, EquiBind’s proportion of high-quality predictions drops further, demonstrating how algorithmic limitations in handling rigid body translations compound with restrictive local energetic landscapes. Co-folding architectures (AF3, Chai-1, Protenix, Boltz-2) maintain an effective ΔMF of 9–13% by leveraging joint receptor-ligand coordinate diffusion to co-adapt local side-chain geometries. Conversely, DynamicBind’s effective ΔMF of 5.8% places it intermediate between rigid docking and full co-folding, reflecting the boundary where induced-fit sampling partially rescues shallow funnels but fails to navigate fragmented energy landscapes. This convergence is not a property of any one architecture. Protenix is an AF3 reproduction, Chai-1 uses ESM-2 embeddings, and Boltz-2 includes affinity and binder-classification heads. The four models differ in training data, objective, and preprocessing. Their convergence on the same effective operating point indicates that they share a common property that dominates other differences between them.

That property is full access to the ligand-induced frustration funnel. The funnel is a feature of the binding landscape: when the ligand binds, the contact set between ligand and protein can reorganize the local landscape into a contiguous minimally frustrated subnetwork that produces a coherent gradient across the interface. A model that can generate and rank multiple candidate poses can sample the resulting basin and converge on its minimum.

While AF3 achieves superior overall docking accuracy compared to DiffDock on the MISATO dataset (57% vs. 29% success rate @ 2 Å; mean QS-score 0.86 vs. 0.71; Table 2), both models produce similar local frustration landscape distributions (ΔMF) (Mann–Whitney U, *p* = 0.08; Cliff’s *δ* = 0.12, negligible). DynamicBind occupies an intermediate position. Its partial induced-fit mechanism allows it to generate candidate poses that partially adapt the receptor, which provides partial funnel access. Its effective ΔMFof 5.8%places it at the Q3/Q4 boundary. The consequence is visible in its quartile distribution because the partial adaptation helps at low-ΔMF sites and is not fully sufficient at high-ΔMF sites. While intrinsic ΔMF represents the physical local frustration quenching of an experimental complex, a model’s effective ΔMF (ΔMF_eff_) denotes its operational benchmark along this energy landscape. For example, EquiBind operates at an effective threshold of ΔMF_eff_ = 1.2% (derived by mapping its mean ligand RMSD to the landscape axis). This indicates that EquiBind’s prediction accuracy drops sharply outside low-frustration regimes, whereas higher-capacity models maintain acceptable pose quality across broader ΔMF values. We argue that the main reason is architectural. EquiBind performs a single-shot regression and produces one deterministic pose per complex, with no sampling and no ranking step. A single-shot model cannot navigate a landscape as it commits to a single point on the landscape, and the accuracy of that point depends entirely on whether the model representation of the pocket and the ligand places the initial guess near the global minimum. When the landscape contains a single dominant minimum (high ΔMF), a well-trained single-shot model has some probability of landing near it. When the landscape contains many shallow minima (low ΔMF), the single-shot model has no mechanism to identify which of the many candidate configurations is preferred, and its prediction remains at whatever point its internal representation happens to generate.

The distinction between DiffDock and EquiBind is therefore not a difference in accuracy but a difference in the class of problem each model can solve. DiffDock solves the same problem as the co-folding models: sample the landscape, rank the samples, return the best. EquiBind solves a different problem: predict a pose directly from the input, with no intermediate landscape exploration. The two problems coincide only when the landscape happens to have a single dominant minimum. At low ΔMF, where the landscape has many minima of comparable depth, the two problems diverge, and the single-shot model has no mechanism to bridge the gap. EquiBind poor performance on MISATO is not an artifact of its architecture; it is the expected behavior of any model that lacks a sampling step, evaluated on a benchmark where the landscape often does not provide a unique solution.

The two axes that organize the behavior of all seven models are therefore properties of the binding landscape topology, sampled by the model. The first axis—funnel accessibility— describes whether the model can engage with the landscape at all. It separates models that generate and rank multiple candidate poses (co-folding cluster, DiffDock, DynamicBind in part) from models that do not (EquiBind). The second axis—exploitation efficiency—describes how precisely the model resolves the minimum once the funnel is accessible (Table 6). It separates the co-folding cluster from DiffDock and DynamicBind, and it explains why AF3 outperforms DiffDock even though both have full funnel access. Neither axis is a property of the model in isolation. Both are defined by the interaction between the model’s sampling capacity and the landscape’s topological organization.

**Table 6.** Two Orthogonal Axes Organize the Behavior of All Seven Models.

| **Model** | **Funnel**<br>**accessible** | **Exploitation**<br>**efficiency** | **High-ΔMF RMSD**<br>**/ success** | **Low-ΔMF RMSD**<br>**/ success** |
| --- | --- | --- | --- | --- |
| AF3 | Yes | High | 2.6 Å / 74% | 5.8 Å / 24% |
| Chai-1 | Yes | High | 2.7 Å / 72% | 6.0 Å / 22% |
| Protenix | Yes | High | 2.7 Å / 71% | 6.1 Å / 21% |
| Boltz-2 | Yes | High | 2.9 Å / 68% | 6.2 Å / 20% |
| DynamicBind | Partial | Moderate | 3.6 Å / 55% | 6.7 Å / 18% |
| DiffDock | Yes | High | 4.4 Å / 38% | 7.2 Å / 14% |
| EquiBind | No | None | 6.2 Å / 12% | 8.4 Å / 4% |

\*High- $\Delta$ MF and low-Δ MF refer to the quartiles defined in Table 3. Success defined as ligand RMSD < 2.0 Å.

The network analysis provides an independent confirmation of this picture. Frustration-weighted protein structure networks, in which shortest paths preferentially traverse minimally frustrated regions, show that the same landscape organization that determines prediction accuracy also determines the position of the binding site in the protein communication architecture. PSN analysis demonstrates that energy landscape funneling correlates directly with global signal propagation topology (Tables S5,S6). Orthosteric pockets are situated on key inter-domain communication pathways characterized by high Shortest-Path Betweenness Centrality (SBC = 0.081 ± 0.019) and elevated relative centrality (*z*-RCA = +2.14± 0.61). In contrast, allosteric and cryptic pockets exhibit significantly lower centralities (SBC = 0.029 ± 0.013, *z*-RCA = +0.38 ± 0.42), placing them on peripheral network clusters. Pockets with low network centrality and low frustration quenching magnitude consistently exhibit decoupling between contact topology (QS-score) and metric coordinate convergence (RMSD) across all evaluated architectures. Across the original orthosteric benchmark, the original allosteric benchmark, and the non-redundant PLA set, the same separation emerges (Table S5). Orthosteric binding sites have a mean normalized SBC of 0.081 ± 0.019, whereas allosteric sites have 0.029 ± 0.013 (*p* < 0.001, Cliff’s *δ* = 0.74). Orthosteric *z*-RCA averages +2.14± 0.61, whereas allosteric *z*-RCA averages +0.38 ± 0.42 (*p* < 0.001, Cliff’s *δ* = 0.81). The fraction of shortest paths through each binding-site residue composed entirely of minimally frustrated edges is 0.68 ± 0.12 for orthosteric sites and 0.22 ± 0.09 for allosteric sites (*p* < 0.001, Cliff’s *δ* = 0.79). The coupling between network topology and frustration was quantified via Spearman correlation between binding-site SBC and ΔMF. On the original orthosteric benchmark, *r*_*s*_ = 0.62 (*p*< 0.001); on the original allosteric benchmark, *r*_*s*_ = 0.14 (*p* = 0.18). This dissociation indicates that the relationship between centrality and quenching is a property of sites that quench and does not hold where quenching does not occur. On MISATO, the correlation is strong across the full dataset (*r*_*s*_ = 0.58, *p*< 0.001) and stronger at the high-ΔMF end (*r*_*s*_ = 0.67 for the top quartiles) than at the low-ΔMF end (*r*_*s*_ = 0.21 for the bottom quartiles), confirming that the topology–frustration coupling tracks the ΔMF spectrum (Table S6). The network analysis therefore extends the frustration findings from a per-residue energetic picture to a global communication-architecture picture, demonstrating that energetic funneling and network topology describe two manifestations of the same underlying physical gradient.

The results suggest that the frustration–predictability relationship is paradigm-independent in outcome but architecture-dependent in accessibility, and the architectural property that determines accessibility is sampling capacity, not the co-folding mechanism. Based on our findings, we argue that the topology–frustration coupling is a general property of protein–ligand binding that generalizes across curated and unbiased datasets, across static and dynamic views of structure, and across co-folding and docking paradigms.

## Discussion

The results establish three empirical relationships that together point to a single physical interpretation. The first is between prediction accuracy and frustration quenching: across all seven models and both benchmark types, ligand RMSD, QS-score, LDDT-PLI, and success rate vary monotonically with ΔMF, with no evident threshold. The second is between frustration quenching and binding-site network centrality: orthosteric sites occupy high-betweenness positions within contiguous minimally frustrated pathways, allosteric sites occupy low-betweenness positions within fragmented neutral regions, and the two descriptors together predict AI success better than either alone. The third is between frustration quenching and sampling capacity: models that generate and rank multiple candidate poses reproduce the gradient, while a single-shot regression model does not access it at any quenching magnitude.

These relationships are consistent with a funnel-based interpretation of binding-site difficulty. Geometric convergence in ligand placement requires a directional gradient in the local energy landscape. Ligand binding can generate such a gradient by resolving strain at the binding site, converting a frustrated region into a contiguous minimally frustrated subnetwork. Sites that undergo this reorganization present a steeper funnel and a more defined low-energy pose; sites that do not retain a flatter landscape with multiple comparably stable configurations, and no unique solution exists for a model to recover. Frustration quenching measures the magnitude of this reorganization directly from apo and holo structures. In this view, the allosteric blind spot is a property of the binding site expressed through its effect on model behavior, rather than a property of the model alone. The interpretation is consistent with the data but remains a hypothesis. The analysis is correlative and does not establish that frustration quenching causes the performance gradient; causal testing would require controlled mutation of the energetic organization of the binding site followed by re-evaluation of prediction accuracy.

The metric hierarchy observed at allosteric sites is consistent with the same picture. Contact recovery, measured by QS-score at a 4.5 Å cutoff, is partially preserved. Local interaction geometry, measured by LDDT-PLI, is partially preserved but degraded. Global pose geometry, measured by symmetry-corrected ligand RMSD, is lost. The ordering follows from the spatial extent of the minimally frustrated subnetwork: a contiguous subnetwork spanning the pocket can support a gradient that specifies both pocket location and ligand orientation within it, whereas isolated minimally frustrated patches anchor specific residue–ligand interactions without producing a gradient across the pocket, leaving the global arrangement underdetermined. The deficit is concentrated at the scale of global pose geometry rather than at the scale of contact identity or local interaction geometry.

The architectural comparison isolates sampling capacity as the discriminating variable among the models examined. DiffDock, a docking model with no co-folding machinery, reproduces the co-folding gradient because it generates and ranks multiple candidate poses. EquiBind, which produces a single pose in one forward pass, does not access the gradient at any quenching magnitude. DynamicBind, whose partial induced-fit mechanism provides only partial access, occupies an intermediate position. Co-folding is therefore not the relevant property; the capacity to generate and rank multiple candidate poses is. This inference is drawn from a limited set of architectures and would benefit from testing on additional models. A related limit is that at the lowest-quenching sites, additional sampling alone may not recover a unique solution, because the landscape does not contain one; improvements at those sites would have to come from elsewhere.

Several limitations bound these conclusions. The general-purpose benchmark samples a wide range of frustration magnitudes but does not provide functional labels, so the analysis cannot address the classification of sites into functional categories. The structural coverage analysis identifies memorization as a contributor to allosteric accuracy but does not separate it from energy-landscape reasoning; a direct test would be to occlude the orthosteric site during prediction and re-evaluate allosteric performance. Dynamic cross-correlation and free-energy landscape calculations from MD simulations would provide an independent probe of funnel depth and barrier heights.

The principal implication of this work is that prediction accuracy can be read as a structural reporter of binding-site energetics, not solely as a measure of model quality. Frustration quenching and network centrality are both computable from the unbound structure, before any prediction is attempted, and both carry information about where a given model is likely to succeed or fail.

## Conclusions

AI docking and co-folding models perform well at orthosteric sites and often exhibit markedly reduced accuracy at allosteric sites, and this gap has been treated as a categorical feature of the two site types. Our results indicate that it is better understood as a quantitative relationship between prediction accuracy and the energy landscape organization that can be characterized from structure topology before any prediction is attempted. By extending the analysis to a general-purpose benchmark assembled without curation for binding-site type, by adding two docking models at opposite extremes of the sampling spectrum, and by introducing a metric that bridges global topology and global geometry, we found that prediction accuracy varies systematically with frustration quenching. The study establishes that AI accuracy loss at non-orthosteric sites is not an algorithmic defect of specific architectures, but a quantitative function of local energy landscape topology—specifically frustration quenching magnitude (ΔMF = MF_holo_ − MF_apo_). By comparing DiffDock (generative diffusion) with EquiBind (single-shot regression), we showed that sampling capacity (generating and ranking candidate poses) dictates whether a model can navigate the frustration funnel. DiffDock reproduces the performance profile of top co-folding models, whereas EquiBind fails to converge even at high ΔMF. The most consequential finding is that the orthosteric–allosteric gap is consistent with a quantitative relationship between prediction accuracy and frustration quenching. Rather than an algorithmic defect to be engineered away, it may be a measurable property of the binding site itself. The frustration quenching magnitude ΔMF and the apo highly frustrated enrichment ΔHFapo make that property visible before any prediction is attempted, and they help explain why sites that are neither strongly orthosteric nor fully allosteric occupy a substantial and consequential part of the landscape. Prior work described a decoupling between contact topology and geometric precision at allosteric sites. Our analysis resolves this decoupling into three tiers corresponding to different spatial scales. Contact recovery, as measured by QS-score with a 4.5 Å cutoff, is partially preserved. Local interaction geometry, as measured by LDDT-PLI, is partially preserved but degraded. Global pose geometry, as measured by symmetry-corrected ligand RMSD, is lost. This ordering is consistent with the spatial organization of frustration: an extended, contiguous minimally frustrated subnetwork can generate a directional gradient spanning the pocket, whereas isolated minimally frustrated patches can anchor specific interactions but do not generate such a gradient. The comparison of seven models suggests that sampling capacity—the ability to generate and rank multiple candidate poses on a frustration-shaped landscape—is more consequential than the co-folding mechanism itself. A docking model with sufficient sampling capacity reproduces the co-folding gradient, while a single-shot regression model does not, even at the highest-quenching sites. This observation reframes the design objective for future methods while also defining its limit: at the lowest-quenching sites, additional sampling alone may not recover a unique solution, because the landscape may not contain one.

Taken together, these findings recast prediction accuracy as a quantitative reporter of energy landscape architecture rather than a simple measure of model quality. Once accuracy is understood in those terms, the allosteric blind spot may become a diagnostic rather than a deficiency, and the two summary metrics introduced here may serve as a practical tool for evaluating and guiding the next generation of energy-aware models. We emphasize that the analysis is correlative, that the mechanistic interpretation remains a hypothesis, and that causal testing through controlled mutation experiments would provide a more decisive test of the framework.

## Author Contributions

Lucas Turano: Software, Validation, Visualization, Investigation, Data Curation, Writing— Original draft preparation, Brandon Foley: Software, Validation, Visualization, Investigation, Data Curation, Writing—Original draft preparation, Ryan Dennis: Software, Validation, Visualization, Investigation, Data Curation, Will Gatlin: Software, Validation, Visualization, Investigation, Data Curation, Writing—original draft preparation, Gennady Verkhivker: Conceptualization, Methodology, Software, Validation, Data curation, Writing-Original draft preparation, Visualization, Investigation, Supervision, Writing-Reviewing and Editing.

## Data Availability Statement

All unprocessed data and original code that the paper reports are freely available in an online repository that meets the criteria for digital longevity, implementation of FAIR standards, and community support. Data for this article, including description of data types, all data, software and scripts are freely available at the GitHub repositories https://github.com/Verkhivker-Research-Group/AI-Co-folding-Allostery-Benchmark/tree/main; https://github.com/Verkhivker-Research-Group/AI-CoFolding-Frustration-Network-Benchmark. All scripts required to reproduce the predictions reported here are publicly available in the GitHub repository. The GitHub repository contains processed prediction outputs, evaluation results, figures, and scripts used in the study. Zenodo archives https://zenodo.org/records/18424406 and https://zenodo.org/records/23173134 provide the datasets, processed prediction outputs, evaluation results, and scripts used in the study. Protein and ligand structures are not included in this archive. All structures are publicly available from the RCSB Protein Data Bank (http://www.rcsb.org) and can be automatically downloaded using the provided scripts. The rendering of protein structures was done with UCSF ChimeraX package (https://www.rbvi.ucsf.edu/chimerax/) and PyMOL (https://pymol.org/2/). The data supporting this article have also been included in the Supplementary Information.

## Acknowledgments

This research was funded by the National Institutes of Health under Award 1R01AI181600-01, 5R01AI181600-02 and Subaward 6069-SC24-11 to G.V. G.V. acknowledges support from Schmid College of Science and Technology at Chapman University for providing computing resources at the Keck Center for Science and Engineering.

All authors have approved the submission of this manuscript, and it has been submitted exclusively to Physical Chemistry Chemical Physics.

