## Supporting Tables S1-S6 for "Frustration Quenching and Network Topology of the Energy Landscape as Primary Determinants of Protein–Ligand Binding Pose Prediction by Deep Learning Models"

**Table S1. Oracle vs. average selection performance across datasets for all seven models**

| **Model** | **Dataset** | **Site type** | **Selection** | **Mean ligand RMSD (Å)** | **Success @ 2 Å (%)** |
|---|---|---|---|---|---|
| **AF3** | Original | Orthosteric | Oracle | 2.3 ± 0.8 | 82 |
| **AF3** | Original | Orthosteric | Average | 3.1 ± 1.2 | 48 |
| **AF3** | Original | Allosteric | Oracle | 6.1 ± 1.9 | 29 |
| **AF3** | Original | Allosteric | Average | 7.2 ± 2.4 | 14 |
| **AF3** | PLA | Allosteric | Oracle | 6.0 ± 1.8 | 30 |
| **AF3** | PLA | Allosteric | Average | 7.0 ± 2.3 | 15 |
| **AF3** | MISATO | — | Oracle | 3.6 ± 1.4 | 57 |
| **AF3** | MISATO | — | Average | 4.8 ± 1.9 | 32 |
| **Chai-1** | Original | Orthosteric | Oracle | 2.4 ± 0.9 | 80 |
| **Chai-1** | Original | Orthosteric | Average | 3.2 ± 1.3 | 46 |
| **Chai-1** | Original | Allosteric | Oracle | 5.2 ± 1.6 | 31 |
| **Chai-1** | Original | Allosteric | Average | 6.6 ± 2.0 | 16 |
| **Chai-1** | PLA | Allosteric | Oracle | 5.7 ± 1.5 | 32 |

| | | | | | |
|---|---|---|---|---|---|
| **Chai-1** | PLA | Allosteric | Average | 6.6 ± 2.0 | 22 |
| **Chai-1** | MISATO | — | Oracle | 3.9 ± 1.5 | 55 |
| **Chai-1** | MISATO | — | Average | 5.0 ± 2.0 | 30 |
| **Protenix** | Original | Orthosteric | Oracle | 2.5 ± 0.9 | 81 |
| **Protenix** | Original | Orthosteric | Average | 3.3 ± 1.3 | 47 |
| **Protenix** | Original | Allosteric | Oracle | 6.2 ± 2.0 | 26 |
| **Protenix** | Original | Allosteric | Average | 7.3 ± 2.5 | 13 |
| **Protenix** | PLA | Allosteric | Oracle | 6.3 ± 1.8 | 27 |
| **Protenix** | PLA | Allosteric | Average | 7.1 ± 2.3 | 15 |
| **Protenix** | MISATO | — | Oracle | 4.0 ± 1.5 | 54 |
| **Protenix** | MISATO | — | Average | 5.1 ± 2.0 | 29 |
| **Boltz-2** | Original | Orthosteric | Oracle | 3.8 ± 1.4 | 74 |
| **Boltz-2** | Original | Orthosteric | Average | 4.9 ± 1.9 | 38 |
| **Boltz-2** | Original | Allosteric | Oracle | 6.8 ± 2.1 | 23 |
| **Boltz-2** | Original | Allosteric | Average | 7.6 ± 2.6 | 12 |
| **Boltz-2** | PLA | Allosteric | Oracle | 6.9 ± 2.0 | 24 |
| **Boltz-2** | PLA | Allosteric | Average | 7.6 ± 2.4 | 13 |

| | | | | | |
|---|---|---|---|---|---|
| **Boltz-2** | MISATO | — | Oracle | 4.1 ± 1.6 | 53 |
| **Boltz-2** | MISATO | — | Average | 5.2 ± 2.1 | 28 |
| **DynamicBind** | Original | Orthosteric | Oracle | 4.1 ± 1.3 | 72 |
| **DynamicBind** | Original | Orthosteric | Average | 4.7 ± 1.7 | 35 |
| **DynamicBind** | Original | Allosteric | Oracle | 5.2 ± 1.5 | 33 |
| **DynamicBind** | Original | Allosteric | Average | 6.8 ± 2.2 | 16 |
| **DynamicBind** | PLA | Allosteric | Oracle | 4.3 ± 1.4 | 34 |
| **DynamicBind** | PLA | Allosteric | Average | 6.2 ± 1.9 | 23 |
| **DynamicBind** | MISATO | — | Oracle | 4.8 ± 1.7 | 41 |
| **DynamicBind** | MISATO | — | Average | 6.0 ± 2.1 | 22 |
| **DiffDock** | MISATO | — | Oracle | 5.2 ± 1.9 | 29 |
| **DiffDock** | MISATO | — | Average | 6.8 ± 2.4 | 18 |
| **EquiBind** | MISATO | — | Oracle | 7.2 ± 2.5 | 8 |
| **EquiBind** | MISATO | — | Average | 8.9 ± 3.1 | 5 |

*Oracle = best-of-25 prediction (lowest ligand RMSD) for co-folding models, best-of-10 for DiffDock, single prediction for EquiBind. Average = mean over all predictions. Values are mean ± SD across complexes. Success defined as ligand RMSD < 2.0 Å. Original = curated orthosteric (2,275 complexes) and allosteric (503 complexes) benchmarks. PLA = non-redundant Protein–Ligand Allosteric set. MISATO contains 9,407 complexes.

**Table S2. Coverage-stratified allosteric performance (oracle selection)**

| Model | Low coverage (≤5 PDB) | Medium (6–20) | High (21–40) | Ultra-high (>40) |
|---|---|---|---|---|
| **AF3** | 6.8 Å / 22% | 6.0 Å / 28% | 5.2 Å / 35% | 4.2 Å / 45% |
| **Protenix** | 6.9 Å / 20% | 6.1 Å / 26% | 5.3 Å / 33% | 4.3 Å / 43% |
| **Boltz-2** | 7.5 Å / 18% | 6.8 Å / 22% | 5.9 Å / 28% | 4.8 Å / 38% |
| **Chai-1** | 6.5 Å / 25% | 5.6 Å / 30% | 4.8 Å / 38% | 4.0 Å / 48% |
| **DynamicBind** | 6.0 Å / 28% | 5.2 Å / 35% | 4.5 Å / 42% | 3.5 Å / 52% |
| **DiffDock** | 6.4 Å / 22% | 5.8 Å / 27% | 5.0 Å / 33% | 4.4 Å / 41% |
| **EquiBind** | 7.8 Å / 10% | 7.4 Å / 12% | 7.0 Å / 14% | 6.2 Å / 18% |

*Values present mean ligand RMSD / success rate at 2.0 Å. Coverage = number of PDB entries sharing the same UniProt ID and pocket residue composition. Low coverage: n = 78 pockets; medium: n = 62; high: n = 32; ultra-high: n = 10. For MISATO, coverage was computed analogously using the number of homologous structures in the PDB.

**Table S3. Conformational Frustration Distributions**

| Dataset | MFsite,apo (%) | MFfold,apo (%) | HFsite,apo (%) | HFfold,apo (%) | ΔHFapo (%) |
|---|---|---|---|---|---|
| **Curated Orthosteric** | 31.2 ± 5.6 | 48.2 ± 4.1 | 27.8 ± 5.2 | 17.3 ± 2.9 | +10.5 |
| **Curated Allosteric** | 21.3 ± 5.8 | 48.5 ± 4.2 | 7.8 ± 2.6 | 17.3 ± 2.8 | −9.5 |
| **MISATO** | 28.4 ± 6.1 | 47.8 ± 4.2 | 18.9 ± 4.5 | 18.1 ± 2.8 | +0.8 |

*Values denote percentages (mean ± SD) of minimally frustrated (MF) and highly frustrated (HF) residues in the apo binding site (site) and global protein fold (fold). $\Delta HF_{apo} = HF_{site,apo} - HF_{fold,apo}$.

**Table S4. Mutational Frustration Distributions**

| Dataset | MFsite,apo (%) | MFfold,apo (%) | HFsite,apo (%) | HFfold,apo (%) | ΔHFapo (%) |
|---|---|---|---|---|---|
| **Curated Orthosteric** | 45.2 ± 6.1 | 44.8 ± 4.5 | 15.0 ± 3.8 | 15.0 ± 3.1 | 0.0 |
| **Curated Allosteric** | 30.2 ± 5.7 | 28.4 ± 3.8 | 11.7 ± 3.0 | 19.2 ± 3.4 | −7.5 |
| **MISATO** | 33.1 ± 6.2 | 28.5 ± 3.7 | 13.9 ± 4.1 | 19.2 ± 3.2 | −5.3 |

*Values denote percentages (mean ± SD) of minimally frustrated (MF) and highly frustrated (HF) residues in the apo binding site (site) and global protein fold (fold). $\Delta HF_{apo} = HF_{site,apo} - HF_{fold,apo}$

**Table S5. Network topology and frustration coupling across datasets**

| Dataset | Stratum | Mean SBC | Mean Z-RCA | MF-only path fraction |
|---|---|---|---|---|
| **Curated** | Orthosteric | 0.081 ± 0.019 | +2.14 ± 0.61 | 0.71 ± 0.12 |
| **Curated** | Allosteric | 0.029 ± 0.013 | +0.38 ± 0.42 | 0.34 ± 0.15 |
| **MISATO** | High ΔMF | 0.068 ± 0.018 | +1.78 ± 0.55 | 0.61 ± 0.13 |
| **MISATO** | Low ΔMF | 0.035 ± 0.014 | +0.52 ± 0.48 | 0.38 ± 0.14 |

*SBC is shortest-path betweenness centrality (frustration-weighted). Z-RCA is Z-score relative centrality analysis. MF-only path fraction is fraction of shortest paths through the binding site composed entirely of minimally frustrated edges. ΔMF strata correspond to the quartiles defined in Table 3 of the main text. Values are mean ± SD across complexes within each stratum.

**Table S6. Spearman correlation between binding-site SBC and ΔMF**

| Dataset | Stratum | ρ | p-value |
|---|---|---|---|
| **Curated** | Orthosteric | 0.58 | <0.001 |
| **Curated** | Allosteric | 0.14 | 0.09 |
| **MISATO** | Full dataset | 0.54 | <0.001 |
| **MISATO** | High ΔMF (top two quartiles) | 0.61 | <0.001 |
| **MISATO** | Low ΔMF (bottom two quartiles) | 0.12 | 0.08 |

*SBC is shortest-path betweenness centrality (frustration-weighted). ΔMF is frustration quenching magnitude. Spearman correlations computed across all complexes within each stratum.
